# The interplay of supercoiling and thymine dimers in DNA

**DOI:** 10.1101/2021.09.27.461905

**Authors:** Wilber Lim, Ferdinando Randisi, Jonathan P. K. Doye, Ard A. Louis

## Abstract

Thymine dimers are a major mutagenic photoproduct induced by UV radiation. While they have been the subject of extensive theoretical and experimental investigations, questions of how DNA supercoiling affects local defect properties, or, conversely, how the presence of such defects changes global supercoiled structure, are largely unexplored. Here we introduce a model of thymine dimers in the oxDNA forcefield, parametrised by comparison to melting experiments and structural measurements of the thymine dimer induced bend angle. We performed extensive molecular dynamics simulations of double-stranded DNA as a function of external twist and force. Compared to undamaged DNA, the presence of a thymine dimer lowers the supercoiling densities at which plectonemes and bubbles occur. For biologically relevant supercoiling densities and forces, thymine dimers can preferentially segregate to the tips of the plectonemes, where they enhance the probability of a localized tip-bubble. This mechanism increases the probability of highly bent and denatured states at the thymine dimer site, which may facilitate repair enzyme binding. Thymine dimer-induced tip-bubbles also pin plectonemes, which may help repair enzymes to locate damage. We hypothesize that the interplay of supercoiling and local defects plays an important role for a wider set of DNA damage repair systems.

## 1 Introduction

DNA supercoiling is actively maintained by cells at significant energetic cost. It plays an important role in key biological processes such as transcription, regulation, and replication, and facilitates the formation of compact structures needed to pack DNA into cellular compartments [1, 2, 3, 4]. Variation in DNA supercoiling is thought to serve as both a sensor and a relay of physiological status that allows the individual cell to optimise its response to changing environmental factors [5]. The movement of the RNA polymerase along DNA during transcription generates downstream positive supercoiling and upstream negative supercoiling. The negative supercoils that are formed facilitate strand separation in the promoter regions of the genome that enable the formation of an open complex with the RNA polymerase [6, 7, 8]. In order to avoid misfunction, excess helical stress is relieved by specialist proteins such as topoisomerases. Supercoiling is also thought to affect DNA-protein binding rates [9] by reducing the energy cost of associated twisting or bending of the DNA [10].

Because of its biological importance, DNA supercoiling has been extensively studied by theory [11, 12, 13] and by *in-vitro* single-molecule experiments [14, 15, 16]. These investigations have mainly considered homogenous double-stranded DNA. Cellular DNA, however, is highly inhomogeneous. For example, many different kinds of proteins bind to DNA and affect its supercoiled structure [1, 17]. In addition, endogenous and exogenous agents can damage DNA, and these defects may have strong effects on local elastic properties [18]. Much less is known about how such inhomogeneities affect the supercoiled structure of DNA and including them into a description of how supercoiling interacts with processes such as replication, transcription, and DNA damage repair, is a major scientific challenge.

In this paper, we focus on the interaction between supercoiling and inhomogeneities caused by thymine dimers, the most common of several types of mutagenic lesions caused by UV irradiation of DNA. They are perhaps the best known and studied DNA damage system [19]. Thymine dimers are also called cyclobutane pyrimidine dimers (CPDs), because they are formed via a cyclobutane ring that connects the 5,6 positions of any two adjacent pyrimidine bases [19]. While it is possible to form an intrastrand cross-link between any adjacent pyrimidine base (i.e. thymine-thymine, cytosine-thymine, and cytosine-cytosine), thymine-thymine dimerisation is the most common.

Persistent thymine dimers are thought to hinder DNA transcription and replication by stalling the progress of RNA polymerase [20, 21]. This can cause cellular death which manifests, for example, in sunburn. Their presence can also lead to the misreading of the genetic code, causing mutations which in some cases can lead to cancer.

Fortunately, living organisms possess a variety of mechanisms to repair detrimental lesions in their DNA [22]. In fact, sunburn only occurs when UV induced thymine dimers are produced at a rate that overwhelms the repair mechanisms. In organisms such as bacteria and plants, thymine dimers can be repaired via enzymatic photoreactivation. A photolyase enzyme binds to thymine dimers and reverses the damage using the energy of light. Mammalian cells, on the other hand, do not have photolyases to perform the repairs. Instead, thymine dimer repair takes place through a light-independent mechanism known as nucleotide excision repair that excises the region of damaged nucleotides instead of just breaking the bonds of the pyrimidine dimer as is done by photolyase. The general excision repair pathway begins with the recognition of the lesion site by the Xeroderma pigmentosum group C (XPC) protein in complex with RAD23B (also commonly referred to as XPC). There is accumulating evidence to suggest that XPC is a non-discriminatory damage marker that is non-specific to CPDs [23, 24]. Thus, for further confirmation that the lesion is indeed a CPD, the multi-subunit transcription factor TFIIH, including the helicase subunits of XPB and XPD, as well as XPA, are recruited [25, 26]. Once the identity and location of the lesion site is verified, XPC dissociates from the complex, and XPG and XPF enters to make the 3’ incision and 5’ incision around the lesion site respectively. The excised oligodeoxynucleotide has a length of approximately 30 bps [27], and is removed from the DNA together with the TFIIH-XPG complex and XPF [28, 29]. Apart from XPC, RNA polymerase can also serve as an initial damage marker [30]. In this transcription-coupled repair pathway, the RNA polymerase stalls at the lesion site on the transcribed template strand, and, unlike XPC, remains on the strand after the dual incision events by XPF and XPG.

Fundamental to these repair pathways is the question of how repair proteins locate and bind to the DNA lesion [31]. In enzymatic photoreactivation for instance, it is generally accepted that the thymine dimer lesion has to be flipped into the active site of the bound photolyase to have the covalent bond between the two pyrimidine bases removed [32]. Yet, it is unclear how this stage of the repair process is achieved in the first place. Two competing models, namely the passive model and the active “induced fit” model, have been proposed. In the passive model, the thymine dimer base pair has to first flip out of the helix before the enzyme can locate and bind to the lesion site [33]. On the other hand, in the active model, the repair enzyme first binds to the lesion site, kinks the duplex at the lesion site (reported bend angles range from 42° in the DNA-XPC/Rad4 complex [34] to 50° in the DNA-photolyase complex [32]), and then flips the thymine dimer bases into an extrahelical arrangement that is fitted into the enzyme’s active site [35, 36]. The passive model seems to be supported by calorimetric calculations [37] and binding kinetic studies [38] but it remains unclear how the CPD lesion adopts an extrahelical conformation in the first instance, given that crystal structure and NMR analyses of thymine dimer-containing DNA typically show an intra-helical arrangement of CPD bases within the double helix [39, 40]. In these configurations, the free energy barrier for flipping out the thymine dimer bases has been calculated to be between 6 to 7.5 kcal/mol, and which is most likely too big to be surmounted by thermal fluctuations on time-scales needed for repair [41].

Many of the experimental and computational studies above that probe the passive/active nature of the repair mechanism use relatively short DNA strands which means that they may miss the potential impact of supercoiling. For example, supercoiling can significantly affect binding affinities [9]. It may also affect the rate at which repair proteins can locate DNA damage. For example, van den Broek *et al*. [42] showed that supercoiling accelerated EcoRV’s search for its target site. In addition, simulations by Brackley *et al*. [43, 44] suggested that for low binding energies, protein target search is significantly shorter if the target sequence is located on a loop of a DNA rosette instead of in the rosette’s centre.

The big question then is how DNA damage interacts with supercoiling to affect protein binding or damage location by repair enzymes. A first step is to ask how damage affects plectonemes. It was suggested in [45] that the formation of a bubble at the tip of a plectoneme (a tip-bubble) could pin the plectoneme. Sequences high in AT content were found to be more likely to form tip-bubbles, and intriguingly, RNA polymerase has been shown to segregate to plectoneme tips [46], suggesting that there is a co-localization of plectoneme tips with AT rich promotor sequences. Furthermore, it was predicted by simulations [48, 47], theory [49], and, importantly, shown by magnetic tweezer experiments [50], that mismatches can also co-localise with plectoneme tips. Taken together, theses previous results suggests a number of hypotheses for thymine dimers. *Firstly*, that they may also co-localise with plectoneme tips, which can lead to pinning that may enhance the ability of repair enzymes to locate thymine dimer damage. *Secondly*, such co-localisation may lead to tip-bubbles, which can enhance the probability that the thymine dimer base pair is in an extrahelical arrangement and may provide a possible pathway for the passive model.

To test these hypotheses in simulations, we need a method that can access the length scales associated with supercoiling, while also accurately describing the breaking and formation of individual base pairs. Atomistic simulations are too computationally expensive to treat this regime, and more coarse-grained models based on the worm-like chain (WLC) model of DNA cannot capture structural changes at the nucleotide level. We therefore turn to the coarse-grained oxDNA model [51, 52, 53], which reproduces DNA’s basic mechanical properties such as the persistence length and torsional modulus, thermodynamic properties such as melting point [53, 54], the behaviour of DNA under overstretching [55], as well as sequence effects on DNA hybridization [56]. In particular, oxDNA has been shown to faithfully reproduce DNA’s response to external tension and torsion [45], including the formation of plectonemes and bubbles (albeit with transitions between different regimes occurring at slightly higher forces than observed experimentally). Furthermore, it has already been used to study the effects of mismatches on supercoiling [48, 47].

We proceed as follows. First, we introduce a description of thymine dimers into the oxDNA model. Then, we show that plectoneme tip-bubbles reveal an extra-helical arrangement of thymine dimers and that these tip-bubbles strengthen the pinning of the plectoneme. Increasing twist increases the probability that thymine dimer base-pairs denature and supercoiling induced plectoneme tip-bubbles further enhances this effect. Finally, we comment on the robustness of our results to model choice, and discuss some potential biological consequences of our findings.

## 2 MATERIALS AND METHODS

### 2.1 Constructing the oxDNA thymine dimer model

Our strategy for modelling the thymine dimers is consistent with the approach used to develop oxDNA, that is, we use a “top-down” coarse-graining approach where a simplified model is parametrised to reproduce experimental structural and thermodynamic properties. This approach differs from a “bottom-up” approach, where a simpler model is derived by integrating out degrees of freedom from a more complex model. Strengths and weaknesses of these two approaches are discussed for example in [57] and [51].

To demonstrate the feasibility of modelling thymine dimers with oxDNA, we therefore need to ensure that the model can reproduce well-established structural and thermodynamic effects of thymine dimers on DNA. Melting experiments have shown that the presence of a thymine dimer can produce a significant drop in the DNA melting temperature in short oligomers [39, 58, 59, 60]. Thymine dimers are also reported to induce a bend in double-stranded DNA. NMR studies suggest a bend angle *θ*_0_ around 10° [39, 61, 62] while the crystal structure of Park *et al*. [40] and the pioneering gel and electron microscope studies by Husain *et al*. [63] suggest that *θ*_0_ ≈ 30°. Atomistic simulations of DNA with thymine dimers do not provide much extra insight into the precise value of *θ*_0_, with estimates in the literature ranging from approximately 15° to 30° [64, 65, 66].

Estimates of the detailed effects of thymine dimers on the helix geometry are also available. NMR Studies by McAteer *et al*. [39] suggest an increase of approximately 26° in the roll angle between the thymine dimer-containing base pairs and a decrease of approximately 9° in the tilt between the same base pairs (from ∼ −1° in the native DNA, to ∼ −10° in the thymine dimer DNA). The crystal structure of Park *et al*. [40], on the other hand, yields an increase in roll of 22° but did not report the tilt. In both studies, the twist angle between the central thymine base pairs of the CPD is reported to decrease, although the magnitude of the decrease varies, with McAteer *et al*. suggesting a decrease of 15° and Park *et al*. a 9° decrease. Results from atomistic molecular dynamics (MD) simulations also show an increase in roll and a decrease in twist between the thymine dimer base pairs (relative to undamaged base pairs) but no change in tilt for the thymine-dimer base pairs [67].

To create a thymine dimer model we focus on reproducing the change in roll angle, DNA melting temperature, and DNA bend angle. There is reasonable consensus on the first two of these properties, but somewhat less agreement on the latter. Changes to tilt were not explicitly incorporated into the model, in part because the introduction of roll is sufficient to generate a bend in DNA, and in part because the results of [67] suggest little change to the tilt.

In the oxDNA model, the stacking interaction term between two adjacent nucleotides on the same strand depends on their relative distance and respective orientations. The changes we introduced into oxDNA to model thymine dimers are illustrated in Figure 1. We modified the stacking term by introducing an increased roll angle to the thymine dimer-containing nucleotide, similar to that observed experimentally. Specifically, the orientation matrix of the nucleotide in the 3’ direction is rotated by an angle *ϕ* around the base vector of the nucleotide before the interaction potential is computed in a typical MD algorithm or energy minimisation simulation. This introduces the desired roll. To demonstrate that the introduction of a non-zero roll angle can generate a noticeable bending of the DNA, we perform potential energy minimisation using the modified stacking interaction term on a 40-bp DNA duplex. This produced the bent DNA shown in Figure 2. For our thymine dimer model, we chose to adjust the roll in a single base pair by 27.1° and enhance the stacking interaction for the thymine dimer-containing base step by a factor of ten to mimic the covalent bonds between the thymines; this model has a DNA bend angle at the potential energy minimum of 13.6°. We also measured the bend angle in MD simulations of the same duplex at *T* = 300 K and a salt concentration of 0.16 M obtaining a value of 11.3° (see Section S2, Figures S1 and S2 in the Supplementary Data). The thermalized value is slightly smaller than at the minimum because the bend fluctuations are not symmetric around the potential minimum. In our robustness analysis below, we also consider a model of thymine dimers that has a larger bend angle commensurate with that measured by Park *et al*. [40].

**Figure 1:**
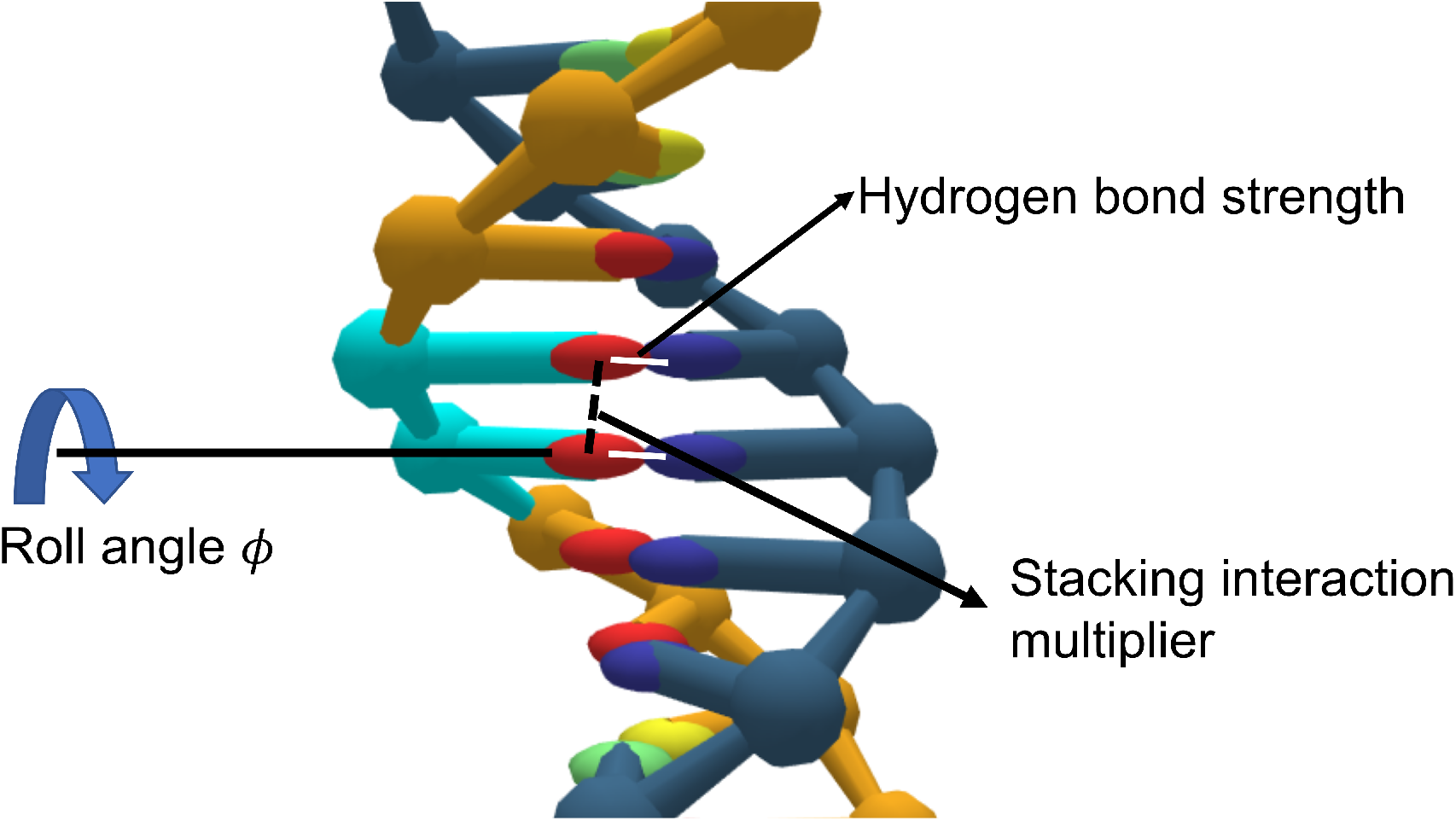
The oxDNA thymine dimer model (highlighted in cyan). Nucleotides are represented by the colours: red (T), blue (A), green (C) and yellow (G). The white lines represent the weakened hydrogen bonding while the strengthened stacking interaction is denoted by a dashed black line. A finite roll of *ϕ* = 27.1° is introduced in our model.

**Figure 2:**
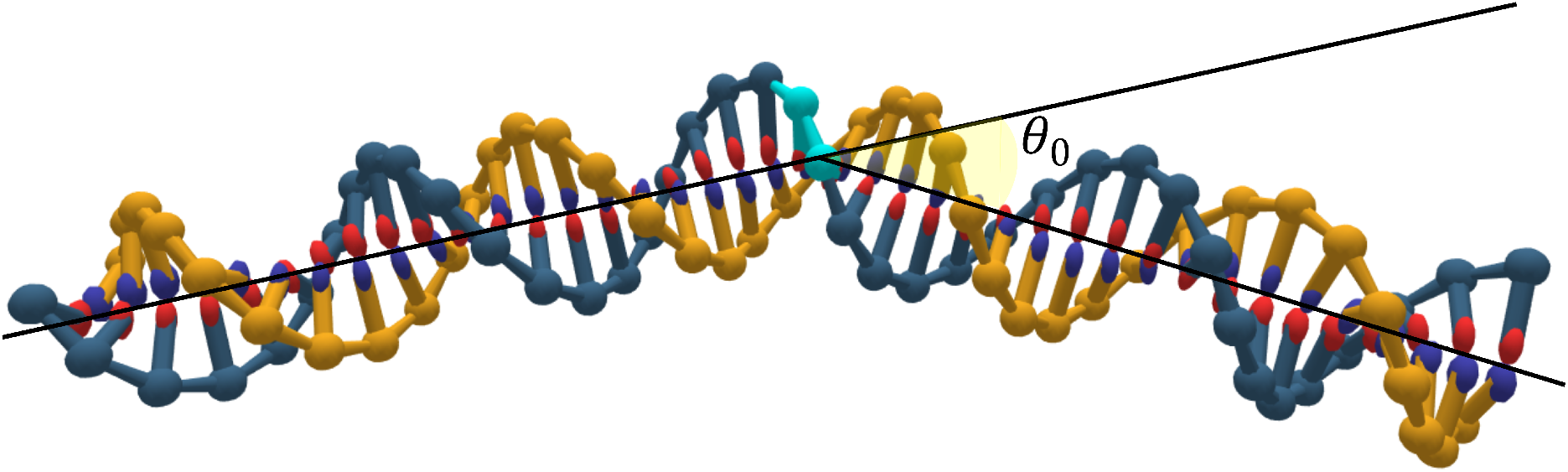
Thymine dimer induced bending: The DNA bend angle *θ*_0_ is measured after energy minimisation is carried out on a 40-bp DNA duplex that contains a thymine dimer (in cyan). The angle *θ*_0_ is defined by first assigning each base pair on the DNA into one of two groups, depending on which side of the thymine dimer it is located. Lines are then fitted through the midpoints of the base pairs of each of the two groups, and used to calculate the bend angle, as in ref. [68].

Next, we used the octamer sequence studied by Kemmink *et al*. [58] as a benchmark for measuring the change in melting temperature induced by the thymine dimer model. These results are similar to the drop in melting temperature found in other studies [39, 59, 60] once the difference in DNA strand concentration used in these studies is accounted for [69, 70]. Umbrella sampling simulations using the Virtual Move Monte Carlo (VMMC) algorithm [71] were carried out (as in [70, 52]) to measure the melting temperature of both undamaged and thymine dimer-containing octamers at the experimental conditions of 6 mM dsDNA concentration and 0.3 M monovalent salt concentration. To ensure that the standard mean error is below 0.2 K, simulations were carried out for at least 5 independent replicas of each system. With just the change to the roll and the stacking detailed above a melting temperature drop is observed, but less than in experiment. To achieve the desired temperature drop, we also reduced the hydrogen bond strengths of the thymine dimer-containing base pairs by 60%. This results in a Δ*T*_*m*_ of 12.3 K, well within the error margins (±3 K) of the experimental value of 13 K.

### 2.2 MD simulation protocols for molecular tweezers geometry

To simulate the effect of supercoiling, we used a similar protocol to that of Matek *et al*. [45], who set up a molecular tweezer geometry using external potential traps. The main difference is our introduction of the thymine dimer defect, and our use of the improved “oxDNA2” [53] model instead of the original oxDNA force-field [51, 52]. A 600-bp DNA duplex, with a randomised sequence with 50% GC content (see Section S3 for the sequence), was simulated with MD at an effective NaCl concentration of 100 mM and *T* = 300 K. 12 base pairs (bps) were attached at each end of the DNA as “handles” to constrain the linking number of the DNA and its spatial position, and to apply the pulling force *F*. The nucleotides in the handle at the “surface” end of the DNA were kept in a fixed position by applying harmonic traps to simulate the presence of a fixed surface to which the DNA is tethered; each nucleotide was subject to an external harmonic potential 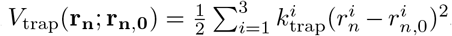,where 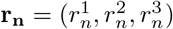 is the centre-of-mass position of the *n*-th nucleotide in the harmonic trap, and the position of the trap **r**_**n**,**0**_ is set to the initial centre-of-mass position of the nucleotide, with 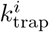 taking on the value of 571 N/m for all the nucleotides in the handle.

At the other end of the DNA duplex, a harmonic potential with 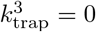 was applied to emulate the freely-moving optical/magnetic bead. This allows the nucleotides to move freely in the direction parallel to the helical axis while remaining constrained in the plane perpendicular to the helical axis. These harmonic traps can also be rotated in the (*x, y*) plane to achieve the desired superhelical density *σ*, here defined as *σ* = (*Lk* −*Lk*_0_)*/Lk*_0_, where *Lk* is the linking number of the supercoiled DNA and *Lk*_0_, the linking number of the relaxed DNA duplex, is 600*/*10.55 ≈ 56.87. For these underwinding/overwinding simulations, a rotation period of 10^7^ MD simulation steps per turn was applied. A constant external force *F/*2 was also applied to the outermost nucleotides of each strand in this handle to model the presence of an external pulling force totalling *F*.

To ensure that the nucleotides do not cross the DNA ends and change the linking number of the system, a repulsive external potential 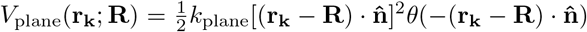 was added to all nucleotides not in the handle. Here, **R** is set to the position of one of the nucleotides in the handle that is furthest from the end of the duplex, *θ* is the Heaviside step function, and 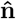 is the normal to the repulsion plane. *k*_plane_ is set to 571 N/m.

MD trajectories were obtained by integrating Newton’s equations of motion with the velocity Verlet algorithm [72] using an integration time step of 6.06 fs. A Langevin-like dynamics thermostat was applied to simulate the impact of the solvent [73]. The diffusion coefficient was chosen such that the diffusive dynamics of individual nucleotides is accelerated by approximately 2 orders of magnitude compared to experiment [74]. This enables the simulation to access longer effective time-scales, accelerating the convergence of the average values of quantities measured during simulations. At least three independent trajectories were used to ensure the reproducibility and consistency of measurements made. A larger than usual number of trajectories were simulated near state boundaries to ensure good sampling of the conformational states. The simulations were performed using the GPU-enabled version of the oxDNA simulation code [75].

To carry out production runs, configurations at the desired superhelical density *σ* and pulling force *F* were extracted from the winding simulations at the appropriate time point. An equilibration time *τ*_eq_ is measured using the decay time of the auto-correlation function of the DNA end-to-end distance. In order to ensure that the observable of interest (i.e. state occupancy, end to-end distance or plectonemic state) is well equilibrated, data was collected for at least 6*τ*_eq_ time steps (between 2 × 10^8^ and 1 × 10^9^ steps). Measurements were made after discarding the first 2*τ*_eq_ time steps.

For simulations involving thymine dimer containing DNA, the same protocol was applied, with the thymine dimer defect placed 110 bps away from the centre of the DNA. Moving the defect away from the centre helps highlight the effect of plectonemic pinning because the constraints on the duplex ends mean that for homogeneous DNA, the plectoneme is most likely to be near the centre of the DNA, an effect that can be quite pronounced for these lengths [45].

## 3 RESULTS

### 3.1 Plectoneme tip bubbles reveal an extra-helical arrangement of thymine dimer

DNA can relieve imposed torsional strain by buckling to form a plectoneme or by denaturing to form bubbles, which have a much lower twist modulus and so can absorb some of the torsional strain. The relaxation pathway that DNA adopts depends on the applied force. For a given superhelical density *σ*, plectonemes are favoured for low forces, and bubbles for larger forces [11, 76, 77].

An intermediate tip-bubble regime where a bubble localizes at the plectoneme tip, was discovered by Matek *et al*. [45] using oxDNA simulations. For this state, the cost of breaking bonds to form the bubble is compensated by several effects. Firstly, the greater ease of bending at the tip leads to a smaller bending cost as well as a reduction Δ*L* of the loop size, which means an additional energy gain of *F* Δ*L*. Moreover, the tip-bubble can absorb some of the torsion just as a normal bubble can. Together, these effects can lead to a tip-bubble regime.

Thymine dimers can enhance plectoneme and tip-bubble formation for two main reasons. Firstly, the intrinsic bending reduces the cost to form an initial loop of the plectoneme. For example, it has also been shown that plectonemes localize at sequences with enhanced curvature [78]. Secondly, the reduced strength of base-pairing lowers the cost of forming a bubble at the thymine-dimer site. Figure 3 depicts such a tip-bubble; the sharp bending at the thymine dimer at the plectoneme tip is typical (quantitative evidence for the thymine dimer’s facilitation of enhanced bending at the plectoneme tip is provided in Section S4 of the Supplementary Data). This configuration also illustrates how this mechanism results in the presence of an exposed unbound thymine dimer at the kinked site of the plectonemic tip-bubble, suggesting the possibility that tip-bubbles may help favour the passive pathway for thymine dimer repair.

**Figure 3:**
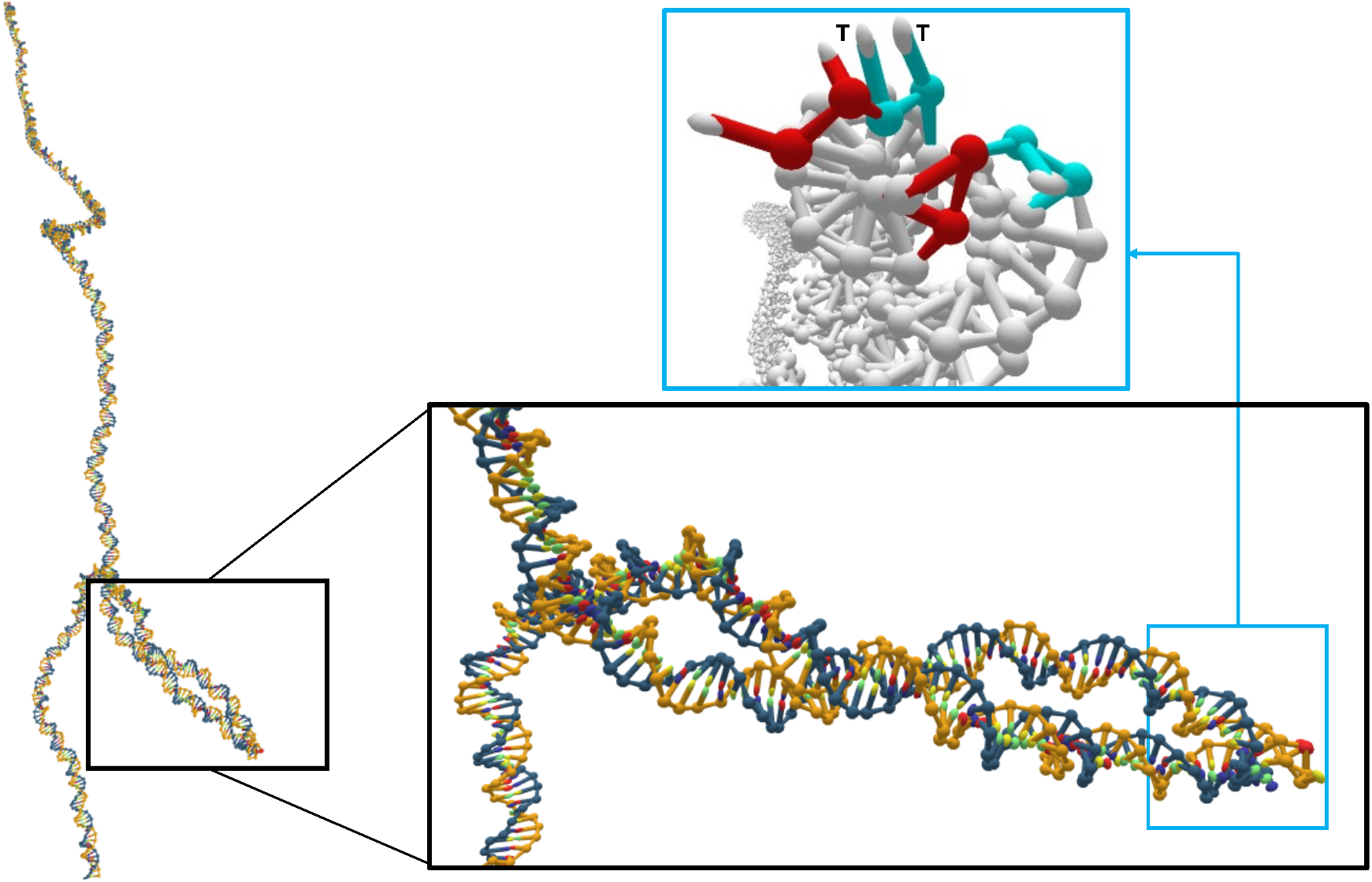
Structure of a 600-bp supercoiled DNA duplex with a thymine dimer 110 bps away from the centre for *F* = 1.5 pN and *σ* = −0.073. The flipped-out thymine dimer base pairs are coloured in cyan (with the thymine bases labelled as T) while the surrounding denatured base pairs are coloured in red (light blue box). The plectoneme possesses an end loop that is highly bent (Section S4, Figures S3-S5), and is pinned by a tip-bubble at the thymine dimer.

To quantify the extent of the tip-bubble regime in thymine dimer-containing DNA, we compare, in Figure 4, the force-torsion state diagrams for undamaged and thymine dimer-containing DNA (quantitative data for the occupancy of the different plectonemic states are given in Figure S6 of the Supplementary Data). As expected, plectonemes start to form at lower *σ* for the damaged DNA than for undamaged DNA (compare *σ* ≈ − 0.038). Tip-bubbles start to form at lower force and *σ* as can be seen, for example by comparing both diagrams at *σ* ≈ − 0.056, and *σ* ≈ − 0.073, for forces of *F* = 1.0 pN and *F* = 1.5 pN, respectively. Such forces and supercoiling densities are typical of values observed *in vivo*.

**Figure 4:**
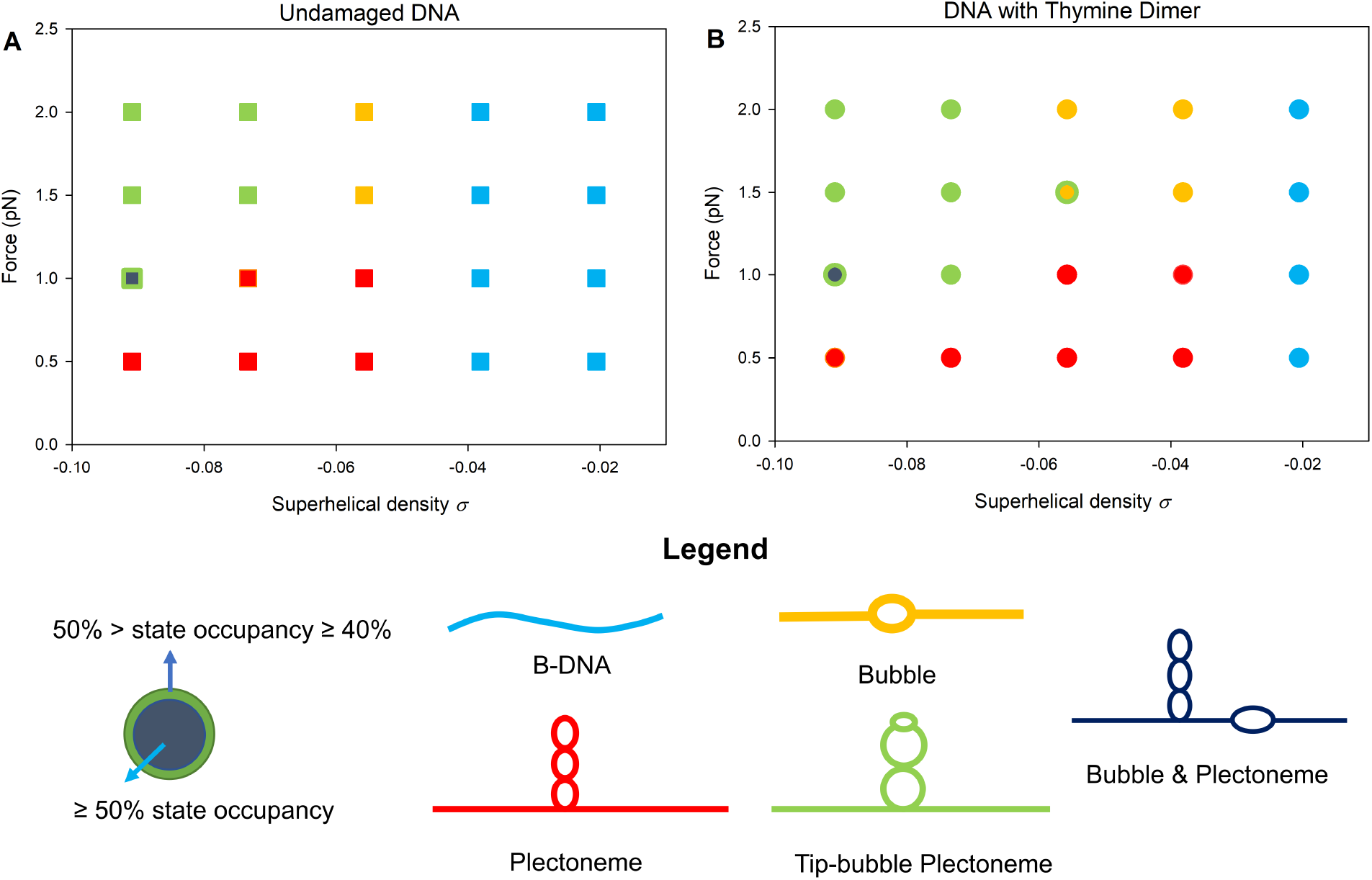
Force-torsion state diagrams for (**A**) undamaged and (**B**) thymine dimer-containing DNA. For each (*σ, F*) point simulated the dominant state is identified. A bubble is defined by the denaturation of at least two consecutive base pairs (a base pair is considered denatured if the binding energy is less than 10% of its maximum possible value). Plectonemes are detected using the algorithm described in the supplementary material of Matek *et al*. [45]. B-DNA is the canonical state where no bubbles or plectonemes are present. A plectoneme is identified as having a tip-bubble, if a kink defect of a few broken base pairs is localised at the tip of the plectoneme (defined here as when the distance between the centre of the bubble and the centre of the plectoneme is less than 20 bps). The dominant states is defined as having a state occupancy ≥ 50%; mixed states with occupancy ≥ 40% but below 50% are also highlighted. At low forces, the presence of a thymine dimer enhances the occupancy of the plectonemic state (relative to the undamaged DNA) at low superhelical densities. The presence of the thymine dimer also increases the tip-bubble plectoneme and bubble state occupancy at low superhelical densities for *F* = 1.5 pN and 2 pN. Quantitative details of the plectoneme state occupancy at the illustrated state points can be found in Figure S6 of the Supplementary Data.

### 3.2 Thymine dimer-induced tip bubbles can pin plectonemes

We also studied the diffusion of plectonemes with and without damage, and representative results at a state point where tip-bubble plectonemes are also observed for the undamaged case can be seen in Figure 5. While tip-bubble plectonemes have previously been shown to be more likely to occur in regions with high AT content [45], we found that the plectoneme in the undamaged DNA is not strongly localised to a specific part of the duplex for the sequence we used here. By contrast, as can be seen for the thymine-dimer containing duplex in Figure 5, the tip-bubbles preferentially form at the site of the damage, and very strongly pin the plectoneme. To verify that the reduced plectonemic diffusivity is directly attributed to the tip-bubble instead of the thymine dimer alone, similar simulations were carried out in the positively supercoiled regime at *F* = 1.5 pN and *σ* = 0.067. Under these circumstances, the plectoneme in the thymine dimer-containing DNA does not have a tip-bubble and possesses much greater diffusivity (see Figure S13 for details).

**Figure 5:**
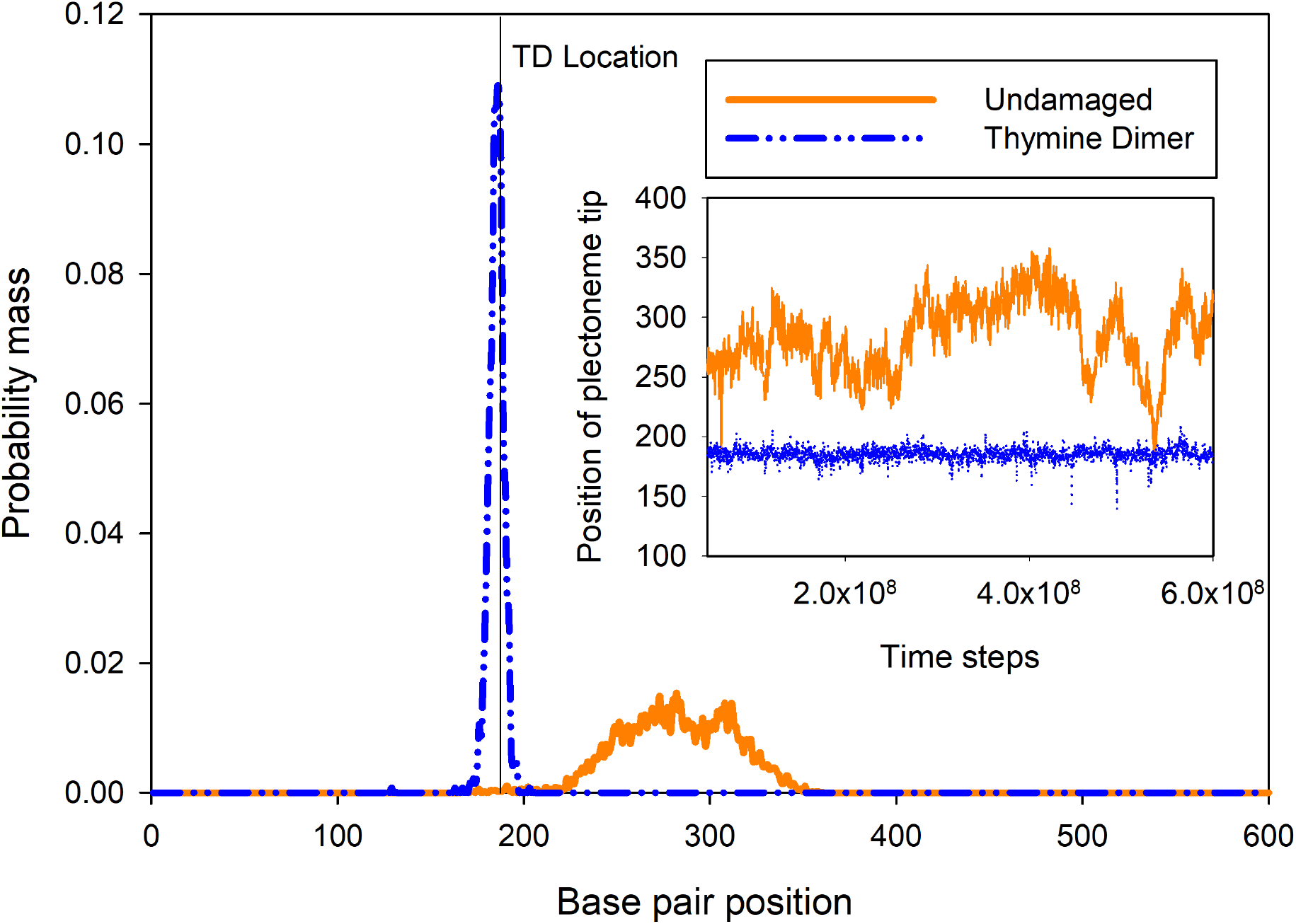
Plectoneme position distributions corresponding to state point *F* = 1.5 pN and *σ* = −0.073. (Inset) The plectoneme position as a function of time. For *F* = 1.5 pN and *σ* = −0.073, we find that for undamaged DNA the plectoneme can diffuse along the duplex, while when a thymine dimer (TD) is present the plectoneme is strongly localised in one place due to the co-localisation of the tip-bubble with the thymine dimer. Using a conversion factor based on the base units of the oxDNA model, the total time of the simulation illustrated in the inset would correspond to 3.6 ms. Because of the reduction in time scale separations inherent to coarse-grained models, using a conversion factor based on diffusional time scales would lead to a much longer simulation time.

### 3.3 Tip bubbles enhance the denaturation probability of thymine dimer base pairs

The stability of the extrahelical arrangement of thymine dimer base pairs can be quantified by measuring the thymine dimer base pairs’ denaturation probability (Figure 6). At low forces and |*σ*|, the thymine dimer base pairs remain largely in an intra-helical conformation for the values of *σ* we studied. By contrast, for the applied tensions of 1.5 pN and 2 pN, the thymine dimer base pairs exhibit a much higher propensity to open up at low superhelical densities relative to its undamaged counterpart. At these forces, we observe that, as a function of increasing superhelical density, first a bubble forms at the site of the thymine dimer, and then subsequently, plectonemic writhe forms with the bubble at the tip, which then causes the thymine dimer base pair to be denatured most of the time.

**Figure 6:**
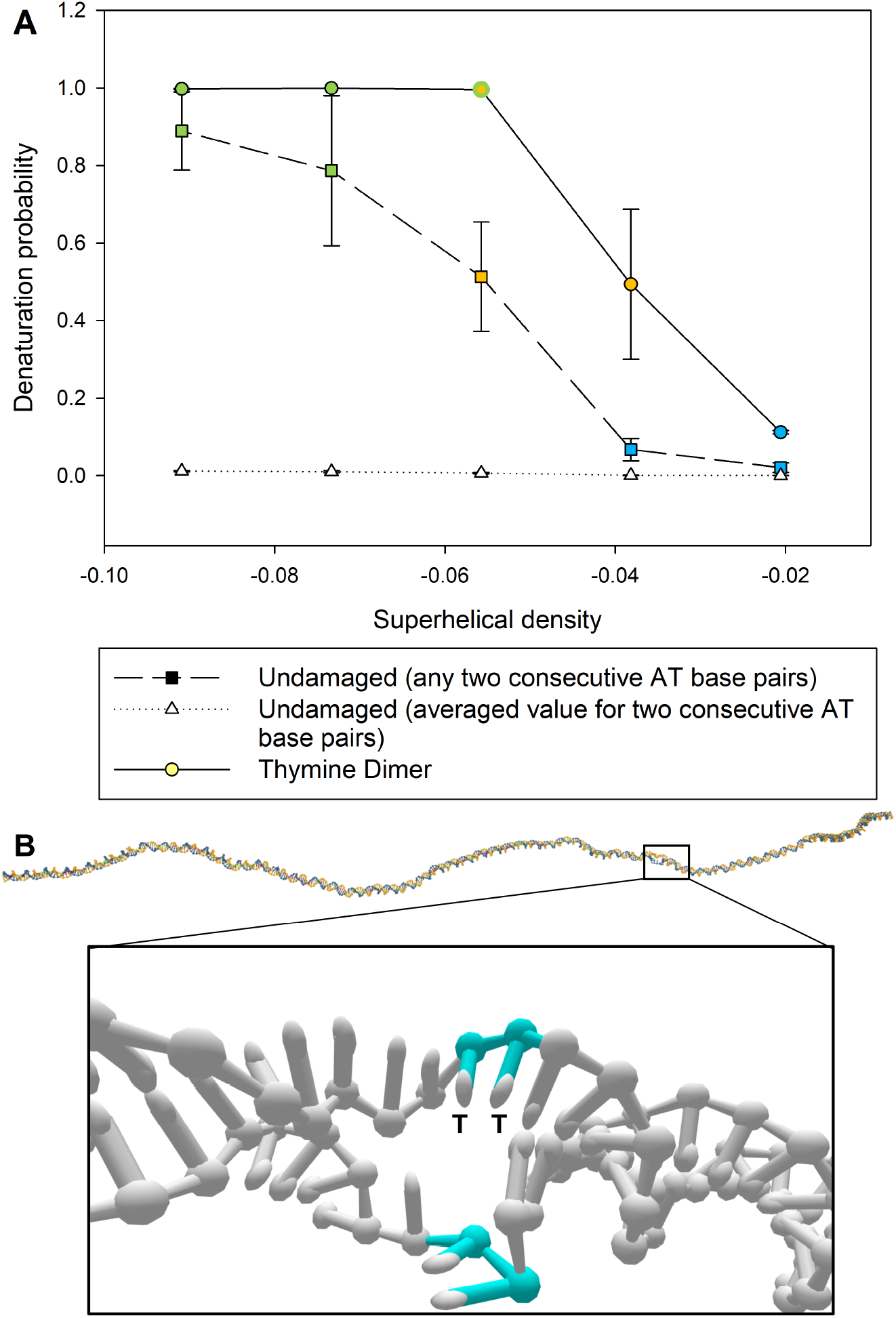
(**A**) Average denaturation probability of the thymine dimer base pairs compared to the probability of *any two* consecutive AT base pairs being open in undamaged DNA at *F* = 1.5 pN. In addition, the dotted line shows the averaged probability that a single set of two consecutive AT base pairs is denatured. The colours in each point correspond to the dominant state occupancy at the given superhelical density (see legend in Figure 4). The thymine dimer base pairs exhibit a higher propensity to denature relative to the undamaged AT base pairs, even at low values of *σ* where no plectoneme or tip-bubbles are observed. At higher forces and superhelical densities, the co-localisation of the plectoneme tip-bubble and thymine dimer causes the thymine dimer to be part of the denaturation bubble most of the time. (**B**) A duplex configuration in the bubble state at *F* = 1.5 pN and *σ* = −0.038. The bubble preferentially nucleates at the thymine dimer site (coloured in cyan with the thymine bases labelled as T). As more turns are imposed, the denatured site becomes the preferred location for plectoneme formation.

### 3.4 Robustness analysis

As an added layer of checks to ensure that the simulation results are not a peculiarity of the parameters we chose for our thymine dimer model, or an artefact of the particular bending angle experiments we chose to replicate, two other models were created. The first additional model, dubbed TD1, had the roll angle *ϕ* adjusted in two adjacent base pairs instead of one and the hydrogen bond strength weakened by 10%. In particular, it produces a larger bending angle, close to that what Park *et al*. [40] observed in their crystal structure. The second model, TD2, has a similar bending angle to the original model, which we will call TD0 here. However, like TD1, it has two base pairs with modified stacking, and like TD0, it has hydrogen-bond weakening of 55%. Table 1 outlines the features of these three models. In all three approaches, there is a linear relation between the DNA bend angle *θ*_0_ and *ϕ* that holds well until *ϕ* ≈ 45° (see Figure S7), thus enabling us to reproduce the different experimental values of DNA bending angle and roll in a consistent manner.

**Table 1:** Features of the three thymine dimer models. Standard errors for the mean bending angles obtained using MD simulations are less than 1.3°

|  | TD0 | TD1 | TD2 |
| --- | --- | --- | --- |
| Roll angle $\phi$ | $27.1^\circ$ | $37.0^\circ$ | $18.3^\circ$ |
| Stacking interaction multiplier | 10 | 1 | 1 |
| Number of base pairs with modified stacking | 1 | 2 | 2 |
| Hydrogen bond weakening | 60% | 10% | 55% |
| DNA bending angle $\theta_0$ (energy minimisation) | $13.6^\circ$ | $27.8^\circ$ | $13.9^\circ$ |
| DNA bending angle $\theta_0$ (MD simulation) | $11.3^\circ$ | $30.7^\circ$ | $13.8^\circ$ |

Increasing the stacking interaction thermodynamically stabilises the DNA duplex, while increasing the roll destabilises it. Thus, models with a different balance between these factors can all adequately reproduce the experimental change in melting temperature. The melting temperatures, which were also computed for TD1 and TD2 using the same VMMC protocols as before are presented in Table 2. The measured Δ*T*_*m*_ for TD1 and TD2 DNA again fall within the error margins of the experimental value.

**Table 2:** Melting temperature *T*_*m*_ of the thymine-dimer containing octamer used in the study by Kemmink *et al*. [58] for the three thymine dimer models compared to undamaged DNA and experiment. The melting temperature drop Δ*T*_*m*_ is within the error margins of experimental results for all three models.

|  | oxDNA |  |  |  | Experiment |  |
| --- | --- | --- | --- | --- | --- | --- |
|  | Undamaged | TD0 | TD1 | TD2 | Undamaged | TD |
| $T_m$ ($^\circ\text{C}$) | 70.4 | 58.1 | 58.0 | 57.0 | 70 | 57 |
| $\Delta T_m$ ($^\circ\text{C}$) | – | 12.3 | 12.4 | 13.4 | – | 13 |

To extract further insights on the different parameter choices of the thymine dimer models, we can also measure the twist at the thymine dimer site. Even though the thymine dimer models were not parametrised to reproduce the decrease in twist angle reported in experiments [39, 40], in relaxed DNA they also all show a similar decrease in the twist angle at the thymine dimer site (Figure S8), albeit of lower magnitude than experiment. In the B-DNA and plectonemic states at *F* = 0.5 pN, the decrease in twist angle at the thymine dimer site increases only slightly (Figures S9A and B). By contrast, in the tip-bubble plectoneme state at higher forces (*F* = 1.5 pN), all three models show significant twist adsorption at the the thymine dimer site, this being largest for the TD0 model (Figure S9C). Similarly, in the bubble state TD0 again shows the most twist adsorption at the thymine dimer (Figure S9D), because of the strong localization at the thymine-dimer site. For TD1 and TD2, twist-induced bubbles can also form at other locations along the DNA strands.

Overwinding/underwinding simulations were also carried out on TD1 and TD2 DNA (Figure S10). We observe a similar increase in state occupancy of plectonemes and tip-bubbles at lower superhelical densities for these models, although there are also some differences in the exact positions of the boundaries between the different states in the three thymine dimer models. With TD1 for instance, bubbles are not the dominant state at *F* = 2 pN and *σ* = −0.038, though its average occupancy at 33% remains higher than its counterpart for the undamaged DNA (23%). On the other hand, TD0 and TD2 exhibit a dominant bubble state at the same point in the force-torsion space. In this particular example, the difference in bubble state occupancy across the three thymine-dimer models can be attributed to the difference in the magnitude of the hydrogen bond strength reduction for these models.

The observation of tip-bubbles at lower |*σ*| than for undamaged DNA gives rise to similar sharply localized plectoneme position distributions for TD1 and TD2-containing plectonemes at *F* = 1.5 pN and *σ* = −0.073 (Figure S11) as found for TD0. With TD1, which is more strongly bent than TD0 or TD2, we also observe plectoneme localization at lower forces and positive *σ* (Figure S12). At *F* = 1.5 pN and *σ* = 0.067, the peak of the probability distribution is approximately half the distribution peak at *F* = 1.5 pN and *σ* = −0.073, and its width (measured at the 50% peak value) is noticeably broader than that of its counterpart at *σ* = −0.073. This indicates that TD1 as a defect on its own constrains the position of a plectoneme somewhat less than a tip-bubble. Under the same sets of *F* and positive *σ*, localization of the plectoneme by TD0 and TD2 is not readily observed. From an energetics standpoint, it requires more bending energy to form a plectonemic end-loop with TD0 or TD2 as compared to TD1 by virtue of their smaller *θ*_0_ at the defect site in the DNA. This in turn makes it less favourable to nucleate a plectoneme at the TD0 or TD2 site.

We can attempt to quantify the favourability of nucleating a plectoneme at the defect site by examining the free energy of plectonemic loop formation. First, we assume that the reduction in the free energy of loop formation at the defect site (relative to the non-defect site) is largely attributed to the bend angle *θ*_0_ induced by the thymine dimerv. <u>Next, using</u> the plectoneme formation energy model by Brutzer *et al*, we can estimate the free energy reduction as 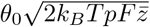,where 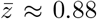 (at *F* = 1.5 pN) is a dimensionless correction term that takes into account writhe fluctuations in B-DNA, and *p* is the bending persistence length of 45 nm [13]. Using the values of *θ*_0_ from Table 1, and *F* = 1.5 pN to ensure a fair comparison with the plectoneme pinning results in Figure S12, we found that the loop formation free energy reduction associated with TD1 is approximately 3.1 *k*_*B*_*T*, while the equivalent values for TD0 and TD2 are 1.1 *k*_*B*_*T* and 1.4 *k*_*B*_*T*, respectively. Consequently, the thermodynamic drive for plectoneme localization at the thymine dimer when a tip-bubble is not present is significantly reduced in the TD0 and TD2 models. Consistent with this, in Figure S13 we observe diffusion of the plectoneme centre away from the defect site for TD0 and TD2 in the positively supercoiled regime, whilst it remains pinned for TD1.

We also compared the denaturation probability of the thymine dimer base pairs for all three models (Figure S14). All show a substantial increase in the denaturation probability of the thymine dimer base pairs at *F* = 1.5 pN when |*σ*| exceeds 0.04. The thymine-dimer denaturation probability is less for TD1 consistent with this model’s lower weakening of base pairing (only 10% less than a normal AT base pair). The smaller energetic advantage of opening a bubble at the thymine dimer rather than elsewhere along the duplex offsets less the entropic cost of localization.

Enhancement to the stacking interaction between thymine bases is present in TD0 but absent in TD1 and TD2 (Table 1). The increased stacking between the thymine bases better mimics the additional covalent linkages in the thymine dimer, and ensures that the thymine bases remain stacked even if the thymine dimer adopts an extrahelical arrangement in the tip-bubble state (Figure 3). This structural effect is clearly absent in TD1 and TD2 models (see section S11, Figures S15-S17 in Supplementary Data for further details). That being said, in our experimentation with various thymine dimer model parameters, we observe that the variation in the stacking interaction multiplier (up to a value of 10) has a relatively small impact on other simulation results (i.e. melting temperature drop, changes in tip-bubble state occupancy, etc.).

## 4 DISCUSSION

Our observations of plectonemic pinning by a thymine dimer are consistent with similar pinning by mismatches, as observed in magnetic tweezer experiments [50] (albeit for positive supercoiling) and also predicted by oxDNA simulations [48, 47] and theory [49]. Taken together, these results suggest that plectonemic pinning may reflect DNA’s structural response to a broader range of possible defects in DNA.

For reasons of computational cost, we have only presented results for a fixed length DNA duplex. To first order, the longer the strand is, the larger the relative entropic cost of localising a plectoneme or a bubble at a damaged site. However, entropic costs typically scale with the log of the length (see for example [49]), so, although the tendency for localization will be reduced at longer lengths, we can still expect qualitatively similar effects from thymine dimers in longer strands. Typical plectonemic loops in bacteria are thought to be on the order of a few thousand to tens of thousands of base-pairs in size [4].

Given that bubble and tip-bubble states are not observed in our simulations at low force (both because the work against the force associated with plectoneme formation is reduced and because the curvature of the plectoneme and its end loop is reduced [79]) one might be tempted to question the relevance of these states to biological systems. First, cellular DNA is subject to transient forces as well as torques by the action of various proteins. Second, in our simulations, we employ a magnetic tweezer-like setup where there are no constraints on the *z* position of the strand end in the twist trap. By contrast, genomic DNA is likely to be much more constrained in its response to superhelical stress by its higher-level organisation. Indeed, small DNA minicircles provide simple examples of the occurrence of tip-bubble and bubble states in the absence of tension due to the circular topology constraining the system’s curvature and leading to an interplay between supercoiling and base-pair opening [80, 81, 82].

Thus, one can instead imagine that the coupling between supercoiling and DNA damage that we have observed might be exploited by biological systems to help achieve certain tasks. One is DNA damage recognition. In our simulations plectoneme localization at a thymine dimer causes it to adopt a configuration that increases its differentiation from normal double-stranded DNA, be it by being more bent or by opening up a denaturation bubble at the plectoneme tip. In the latter case, the thymine dimer is likely to adopt an extrahelical configuration. These more extreme configurations could well allow the DNA damage to be more easily recognised by the relevant protein.

In a crowded living cell, a large number of proteins simultaneously seek to locate specific targets on the chromosomal DNA to carry out their biological functions. Negatively supercoiling could potentially aid the discovery of the thymine dimer (and other defects) by favouring the location of the damage in the relatively easy-to-find plectonemic loop away from the crowded environment at the centre of the chromosome. To explore this hypothesis by simulation, much longer DNA duplexes in much more complex configurations must be treated than the simple geometry that we use here.

Unlike in our simulations where a fixed superhelical density and force is applied, cellular DNA is constantly being subjected to transient local mechanical stresses applied by various proteins. The coupling between their action and the mechanical response of defective DNA might also provide a means to locate DNA damage, such as thymine dimers. For example, in the the twist-open mechanism of damage recognition proposed for the Rad4/XPC nucleotide excision repair complex [83, 24], the requisite repair protein switches between “search” mode (i.e. facilitated diffusion) and “interrogation” mode (i.e. unwinding of the DNA duplex) in its attempt to locate the defect on DNA. One could envisage that a plectoneme is preferentially formed at the defect site when the repair protein interacts with and twists the DNA duplex during the interrogation mode. The structural distortion then serves as a marker for the repair protein, stalling its diffusive motion and enabling it to engage more extensively with the thymine dimer site.

The increased probability of bubble formation at a thymine dimer means that double-stranded DNA is more likely to absorb excess twist at such damage locations. Various biological processes such as transcription and replication are affected by the denaturation profile [84], which might in turn be affected by the presence of a thymine dimer. Such twist absorption effects were indeed proposed over forty years ago [85], and there are examples in the literature of thymine dimers and other damage affecting replication efficiency in plasmids [86, 87], although the effects observed there may be more complex than just the twist absorption effect mentioned above [81]. Moreover, it remains to be understood how excess twist can be partitioned between defect sites and twist buffering protein complexes in vivo such as chromatin fibers in eukaryotic DNA [88].

Recent developments in DNA nanotechnology have shown that user-defined placement of thymine dimers in DNA nanostructures can provide an additional level of structural stability. In this approach UV irradiation is used to generate thymine dimers that link strand ends and bridge strand breaks at crossover sites [89]. Studying how these thymine dimers affect DNA nanostructures is another avenue of research where the oxDNA model, which has been widely used to study such systems [90, 91, 92], can now be applied.

Although we have shown that plectoneme pinning by a thymine dimer is robust to variations in our model, it would of course be desirable to further explore these phenomena with more detailed atomistic simulations [93, 80, 94]. While larger scale writhing effects such as the formation of plectonemes would be very challenging to study with atomistic simulations, it may be possible to use multi-scale techniques to treat the writhe on a coarser level, for example with oxDNA, and to then locally treat the thymine dimer with more fine-grained solvent-free atomistic force-fields.

Finally, the best way to test these hypotheses about the interaction between supercoiling and DNA damage is through experiments. One promising direction is through *in-vitro* single-molecule experiments with magnetic tweezers. These can be used to test, as was done in [50] for mismatches, whether thymine dimers pin plectonemes. Using fluorescent probes, it should also be possible to investigate how imposed force and twist affect the propensity of thymine dimers to denature. It may also be possible to measure the binding constants for key thymine dimer repair proteins as a function of superhelical density, as was done in [9] for example. Demonstrating whether or not these mechanisms are at work and assist the location and repair mechanisms of thymine dimers and other defects *in vivo*, is a significantly more complex challenge.

## 5 CONCLUSION

To conclude, we have demonstrated a working model of thymine dimers in oxDNA that can reproduce known experimental results. There is a certain amount of robustness in the choice of model parameters, constrained by experiments on melting temperatures and bending angles. Within this range of parameters, we achieve consistent results on DNA’s response to twist, namely a preferential co-localization of the thymine dimer with the tips of supercoiled plectonemic structures and the enhanced probability for the thymine dimer to be in a bubble state. This robustness suggests that the results are not a peculiarity of the parameters used in our model, but are instead reflective of a more general interaction between supercoiling and the thymine dimer. By exposing the thymine dimer in a plectoneme tip-bubble, negative supercoiling could play a bigger role than previously thought in facilitating the repair of thymine dimers. Moreover, our results here are suggestive of a much richer set of possible mechanisms that could be driven by the coupling between supercoiling and DNA damage.

## 6 DATA AVAILABILITY

The oxDNA simulation code is available to download at https://sourceforge.net/projects/oxdna/. The GPU implementation of the thymine dimer model can be found in the /contrib/randisi folder of the oxDNA package.

## 7 ACKNOWLEDGEMENTS

The authors acknowledge the use of the University of Oxford Advanced Research Computing (ARC) facility (http://dx.doi.org/10.5281/zenodo.22558) and the resources provided by the Cambridge Service for Data Driven Discovery (CSD3). We thank Christan Matek, Timothy Craggs, Michael Selby, and Lorenzo Rovigatti for discussions on defects and supercoiled DNA, and Troy Lionberger for pointing out ref [46] to us.

