## Supplementary Data for "The interplay of supercoiling and thymine dimers in DNA"

#### S1 oxDNA model modifications for the thymine dimer

##### Stacking and hydrogen bond strength modifiers

Let $\epsilon_{\text{stack}}$ and $\epsilon_{\text{HB}}$ be the stacking and hydrogen bond strength modifier respectively. The stacking and hydrogen bond potentials $V_{\text{stack}}$ and $V_{\text{HB}}$ as defined in Appendix A of reference [1] are now replaced via:

$$\begin{aligned} V_{\text{stack}} &\longrightarrow \epsilon_{\text{stack}} V_{\text{stack}} = \epsilon_{\text{stack}} f_1(r_{\text{stack}}) f_4(\theta_4) f_4(\theta_5) f_4(\theta_6) f_5(\cos(\phi_1)) f_5(\cos(\phi_2)) \\ V_{\text{HB}} &\longrightarrow \epsilon_{\text{HB}} V_{\text{HB}} = \epsilon_{\text{HB}} f_1(r_{\text{bond}}) f_4(\theta_1) f_4(\theta_2) f_4(\theta_3) f_4(\theta_4) f_4(\theta_7) f_4(\theta_8) \end{aligned} \tag{S1}$$

The functional forms and variables are defined in reference [1].

##### Introduction of a positive roll to modify stacking potential

Let $\hat{\mathbf{n}}$ and $\hat{\mathbf{b}}$ be the base normal and backbone-base vector (respectively) as defined in Appendix B of Ref. [1].

Let $\phi$ be the (positive) roll that we introduce to a nucleotide in order to modify $V_{\text{stack}}$. This is achieved by rotating the base normal about the backbone-base unit vector by $-\phi$ (right hand rule):

$$\hat{\mathbf{n}}_{\text{rotated}} = \hat{\mathbf{n}} \cos\phi - \left( \frac{\hat{\mathbf{b}}}{|\hat{\mathbf{b}}|} \times \hat{\mathbf{n}} \right) \sin\phi + \frac{\hat{\mathbf{b}}}{|\hat{\mathbf{b}}|} \left( \frac{\hat{\mathbf{b}}}{|\hat{\mathbf{b}}|} \cdot \hat{\mathbf{n}} \right) (1 - \cos\phi) \tag{S2}$$

The stacking interaction is then the same as given in Ref. [1] but with $\hat{\mathbf{n}}$ replaced by $\hat{\mathbf{n}}_{\text{rotated}}$. The original $\hat{\mathbf{n}}$ is used in all the other oxDNA interaction terms.

In the TD0 model, this modification is applied to the $3'$ nucleotide in the thymine dimer, but in the TD1 and TD2 models it is applied to both thymines. As the value of the bend angle $\theta_0$ varies approximately linearly with the roll angle $\phi$ introduced at the thymine dimer (Fig. S7), it is straightforward to choose the $\phi$ required for a desired bend angle. The effect of the roll modification being applied to both thymines, rather than just one, unsurprisingly leads to a larger bend angle for a given $\phi$ albeit being less than twice as large.

### S2 MD simulations to measure double-stranded DNA (dsDNA) bend angle

MD simulations were carried out at $T = 300\,\mathrm{K}$ and a salt concentration of $0.16\,\mathrm{M}$ on the same 40-bp DNA duplex used in the energy minimisation. The length of the simulation runs were $10^8$ time steps (using an integration time step of $6.06\,\mathrm{fs}$). For each system of interest at least three independent runs were carried out. The dsDNA bend angle reported in the paper is the averaged value obtained from these runs, each with a standard mean error of 1.3° or less.

##### Determining the bend angle

The bend angle's definition is adapted from the one used in energy minimisation simulation (Figure 2 in main paper). Each base pair on the DNA is assigned into one of two groups, depending on which side of the thymine dimer it is located. Lines are then fitted through the base pairs' centres of mass of each of the two groups, and used to calculate the bend angle $\theta_0$. In defining the centre of mass of a base pair, a small offset in the direction of the major groove is introduced to correct for the helicity of the duplex contour [2]. Because fraying of base pairs at the end of the dsDNA was observed in our simulations, the terminal base pairs were not used in line fitting. The thymine dimer and the base pairs adjacent to the dimer are also not included in the line fitting.

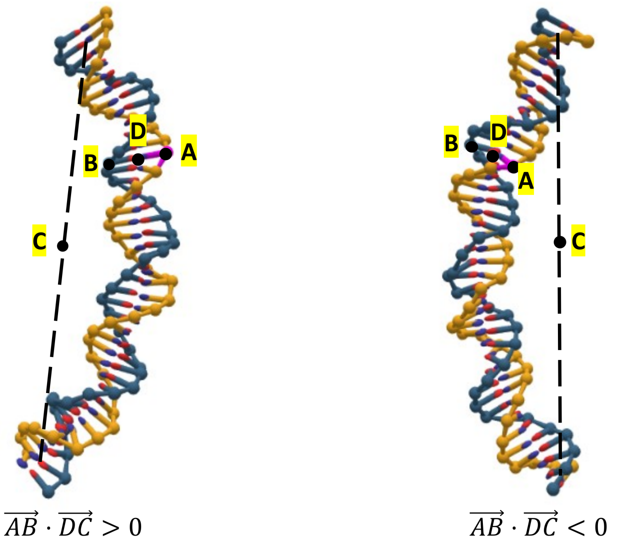


Figure S1: Points A and B are taken to be the centre of mass of off-centre nucleotides, while D is the mid-point of A and B. C is taken to be the centre of mass of the second last base pairs on both ends of the dsDNA.

The sign of the bend angle is determined using the convention outlined in Figure S1. When $\vec{AB} \cdot \vec{DC} > 0$, the angle is taken to be positive. Likewise, when $\vec{AB} \cdot \vec{DC} < 0$, the angle becomes negative. The calculated bend angle is observed to vary with the position of $\vec{AB}$. Thus, we choose to position $\vec{AB}$ at a distance of 7 bps away from the thymine dimer as this gives a maximum calculated bend angle for the damaged DNA. The bending induced by the thymine dimer is roughly towards the major groove. Using this convention, the bend angle of the undamaged dsDNA was found to be $-0.1 \pm 1.3°$

while the bend angle of the thymine dimer (TD0)-containing dsDNA was calculated to be $11.3 \pm 0.3°$.

The standard deviation $\sigma_\theta = \sqrt{\langle\theta_0^2\rangle - \langle\theta_0\rangle^2}$ of the bend angle fluctuations was also measured (Fig. S2), but we did not find any significant difference between the standard deviation of the bend angle fluctuations in undamaged dsDNA ($\sigma_\theta = 28.5 \pm 0.1°$) and that of the thymine dimer (TD0)-containing dsDNA ($\sigma_\theta = 31.0 \pm 0.1°$). The full probability distributions for $\theta$ are illustrated in Figure S2.

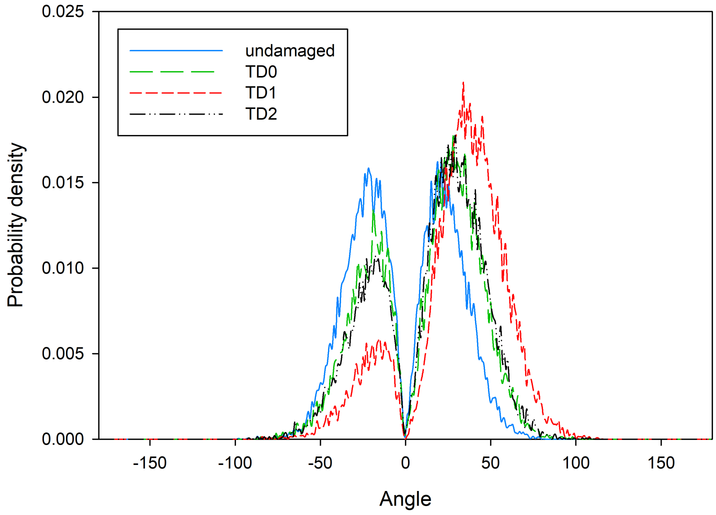


Figure S2: Probability distribution of bend angles for the undamaged and thymine dimer-containing dsDNA. A bin width of 1° was used. Standard deviations for the bend angle fluctuations in TD0, TD1 and TD2 are respectively 31.0±0.1°, 28.7±0.6° and 30.7±0.6°.

#### S3 DNA sequence used in the simulations

The position of the thymine dimer is indicated by circumflexes on the relevant thymines.

3’-AAGCCGGCCACGCGCTGTTCGGCCGGCATTTACACCAAACAGATTGTCAGGA
ACTGTATCTTAGCTCCCGTTTCTGGACTTACGTGCTCAGGTGGAAAACCTGGTA
GACCTAACCGGACTCTACGCTTATGCCAGGGCAATCTTCAACGATAGCTTTCCC
TAGGCGGCTATAATCCACCCTACCTGTTT̂T̂TGGTTCGAGGTCACGGATTCAAGA
TCTTGTCGGATACGGCGCGCGAGTCCACGAGCCCTCGTTAGTTGTCCTCGCGG
GAATAGCCGACGATTAATCTGAAACCGCCGTCGGGTCTCGTGATAGCTACATCG
GTGGATAAACTTCCTAGGCGCTAAGTCCACACACGCGCGCCCTATAGATGAGGA
CCACCAAGTCGAACCGTGCGAATCGTCCGAGAATTCCCCCAACGACAAAATTTC
GTGCGAAACGGTTGCTAAAGGAATAAATGCGTCAGAGCGATTTAGGGGACCCC
GCCCGTGGTTTGGTAGGCTGTTGATTTCTTTCAGACTGTAAGGACCGGCCCAG
CTCTGCGAAGTGTGCTCGCTAGCCGGCACCGAAGACATACATTAAATGGACACC
CGGGAAGACGA-5’

### S4 End-loop angle of tip-bubble plectonemes

We measure the end-loop angle of tip-bubble plectonemes in undamaged and thymine dimer-containing dsDNA, following closely the definition of end-loop angle used by Desai *et al.* in their paper [3]. Taking the centre of the plectoneme to be $i$, three blocks of 5 bps are defined along the end-loop: $i-12$ to $i-8$ bp (block 1), $i-2$ to $i+2$ bp (block 2), and $i+8$ to $i+12$ bp (block 3). The end-loop angle is then obtained from the dot product of the vectors connecting the centre of masses of these three blocks (i.e. block 1 and 2, as well as block 3 and 2). Figure S3 illustrates this definition.

Since the plectoneme detection algorithm by Matek *et al.* [4] does not always pinpoint the exact location of the plectoneme tip, the centre of the plectoneme $i$ is varied by 15 bps on each side. The smallest value of the angle that we obtain from this variation is then set as the end-loop angle (i.e. we are effectively defining the tip of the end-loop as the point with strongest bending).

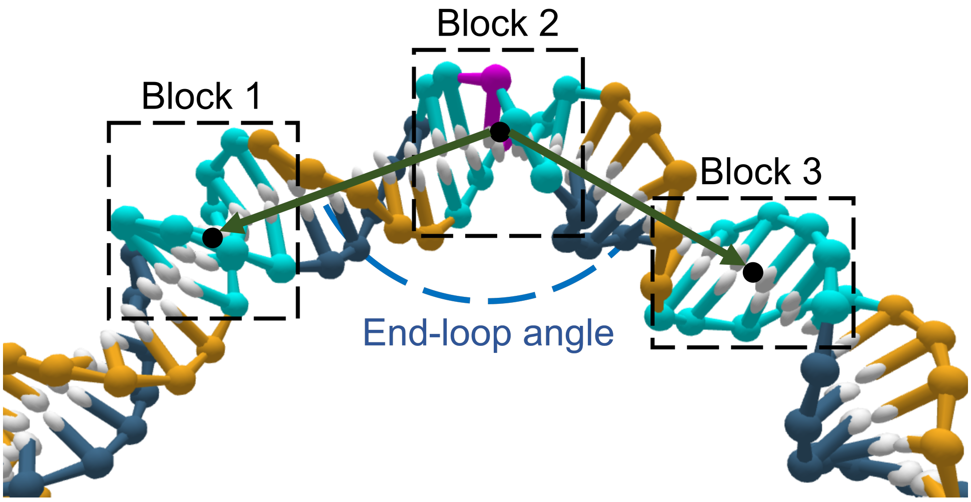


Figure S3: Definition of the end-loop angle. In this illustration, the tip of the plectoneme is taken to be the base pair coloured in purple (base pair $i$). From there, three blocks of 5 bps (coloured in cyan) are defined: $i-12$ to $i-8$ bp (block 1), $i-2$ to $i+2$ bp (block 2), and $i+8$ to $i+12$ bp (block 3). The end-loop angle is then defined as the angle between the vectors connecting the centres of mass of blocks 1 and 2, and blocks 3 and 2.

In Fig. S4 we show the results for the end-loop angle in a regime where the tip-bubble plectoneme is the dominant state (Fig. 4). The end-loop is much more strongly bent when the thymine dimer is present (see Fig. S5 for an illustration).

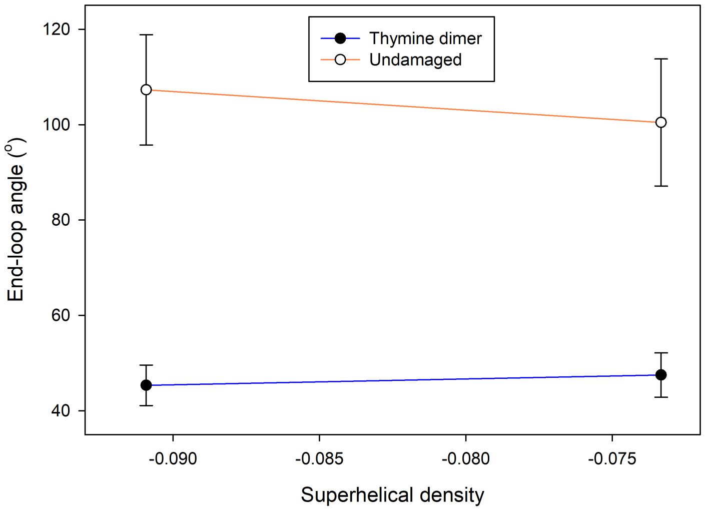


Figure S4: The end-loop angle of the tip-bubble plectoneme is measured at $F = 1.5\,\text{pN}$ at two superhelical densities $\sigma$ where the tip-bubble plectoneme is the dominant state in both undamaged and thymine dimer-containing dsDNA. At $\sigma \approx -0.073$ and $F = 1.5$ pN, the tip-bubble plectoneme occupancy rates are 0.65 and 0.93 for the undamaged and thymine dimer-containing dsDNA respectively. At $\sigma \approx -0.091$ for the same value of $F$, the occupancies become 0.71 and 0.99 for the undamaged and thymine dimer-containing dsDNA respectively. Error bars represent the standard error of the mean.

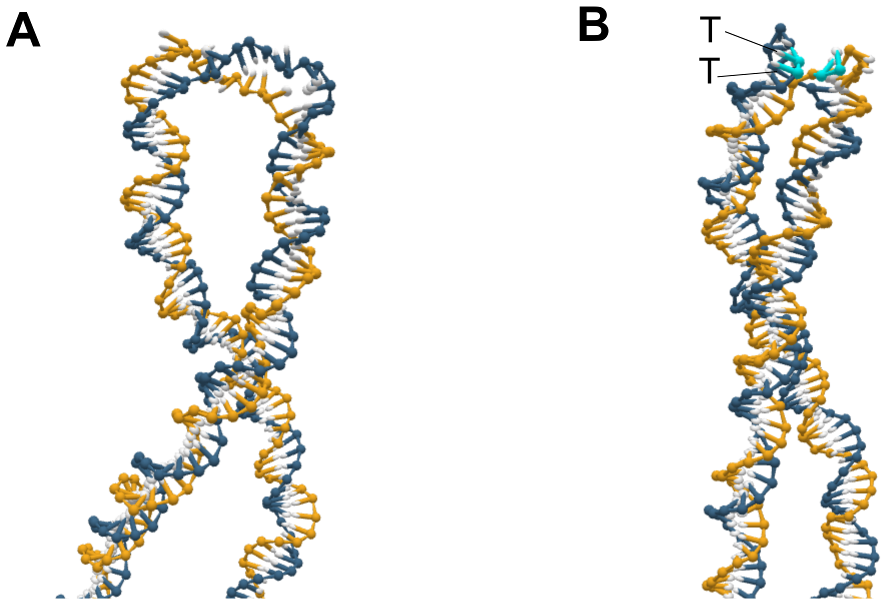


Figure S5: End-loop of a plectoneme with a tip-bubble for (**A**) undamaged dsDNA and (**B**) dsDNA containing a thymine dimer. Both are from simulations at $\sigma \approx -0.091$ and $F = 1.5$ pN. The thymine dimer is highlighted in cyan and the thymine bases are labelled with 'T's.

### S5 Occupancy of the plectonemic state

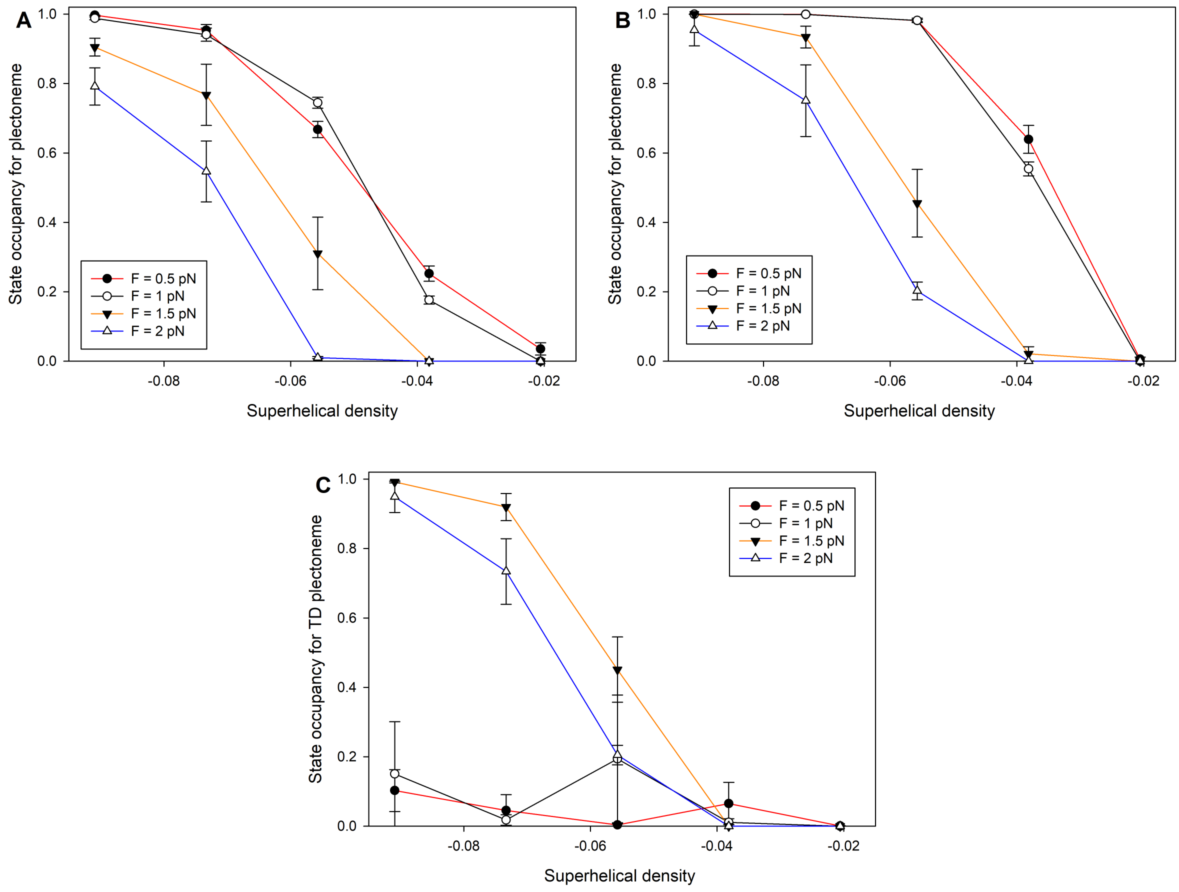


Figure S6: State occupancy for the plectonemic state in (**A**) undamaged dsDNA and (**B**) thymine dimer-containing dsDNA. Error bars represent standard errors of the mean. At $F = 1.5$ pN and 2.0 pN, the occupancy of the plectonemic state in both undamaged and thymine dimer-containing dsDNA is strongly correlated with the state occupancy of the tip-bubble plectoneme. In (**C**), the TD plectoneme is defined as a plectoneme with a thymine dimer (TD) within 20 bps of its tip (based on the plectoneme detection algorithm by Matek *et al.* [4]). The thymine dimer is shown to be strongly localised at the plectoneme tip when the dsDNA is in the tip-bubble plectonemic state at $F = 1.5$ pN and 2.0 pN.

### S6 Relation between dsDNA bending and roll in the TD models

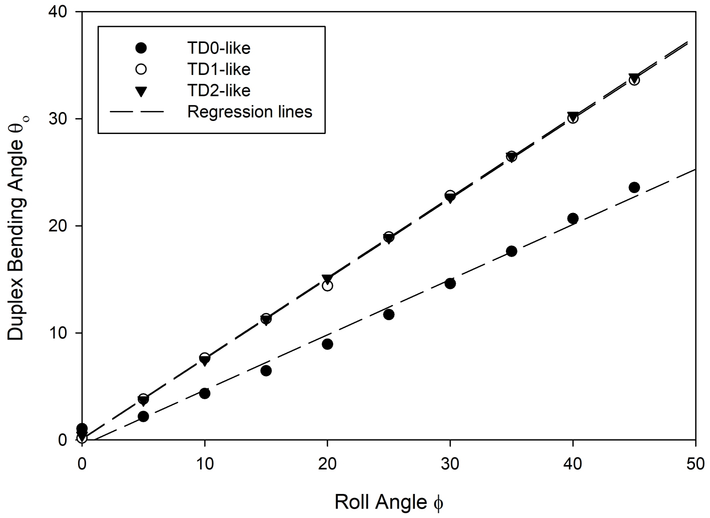


Figure S7: Energy minimisation is carried out on a 40-bp DNA duplex to generate the plot of $\theta_0$ as a function of $\phi$. $\theta_0$ increases approximately linearly with $\phi$ until $\phi \approx 45°$. Beyond that the stacking breaks and the dsDNA bending angle decreases significantly.

### S7 Twist Angles at the thymine dimer site

The definition of twist angle follows from the approach taken by Skoruppa *et al.* [5]. A rotation vector $\boldsymbol{\Theta}^{(i)}$ represents the rotation of the orthonormal triad $\{\hat{\mathbf{e}}_1(i), \hat{\mathbf{e}}_2(i), \hat{\mathbf{e}}_3(i)\}$ associated with base pair $i$ onto the adjacent triad $\{\hat{\mathbf{e}}_1(i+1), \hat{\mathbf{e}}_2(i+1), \hat{\mathbf{e}}_3(i+1)\}$, where $\hat{\mathbf{e}}_1$ is oriented towards the major groove, $\hat{\mathbf{e}}_2$ is along the direction of base pairing and $\hat{\mathbf{e}}_3$ is along the DNA contour. The twist angle between these two base pairs is given by the third component of the rotation vector $\Theta_3$. In all our twist angle calculations in this paper, $\hat{\mathbf{e}}_3$ is defined using "Triad III" that is described in the supplementary materials of Ref. [5].

Twist angle measurements on the same thymine dimer-containing 40-bp duplex used in the bend angle measurements in Section S2 are presented in Figure S8 as a reference point for a system that is subject to no torsional stress. There is a model-specific pattern of slightly reduced twist angles close to the thymine dimer. Experiments also suggest a reduction in the twist angle at the thymine dimer. Fig. S9 reports the twist angle measurements for the thymine dimer-containing dsDNA in different states (i.e. twisted B-DNA, plectoneme, bubble, tip-bubble plectoneme). In twisted B-DNA and plectonemic DNA when the plectoneme is not localized at the thymine dimer the pattern of twist angle reduction at the thymine dimer is very similar to the unstressed reference albeit with a very small increase in the twist angle reduction. However, when a tip-bubble or bubble is localized at the thymine dimer, the picture is completely different; there is clear absorption of twist at the site of thymine dimer, even leading to large negative left-handed twists close to the thymine dimer.

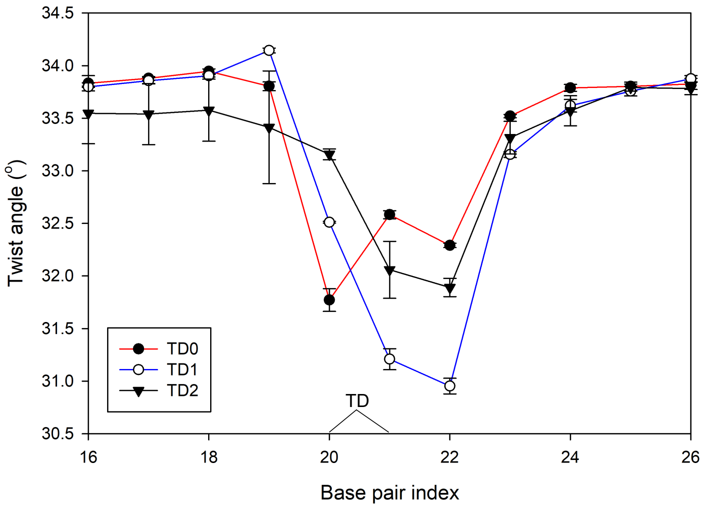


Figure S8: Twist angles measurements for the thymine dimer-containing 40-bp duplex used in Section S2. The value indicated at base pair index $i$ represents the twist angle between base pair $i$ and base pair $i-1$. Error bars represent standard errors of the mean. The thymine dimer is located at positions 20 and 21 and is marked by "TD" in the plot.

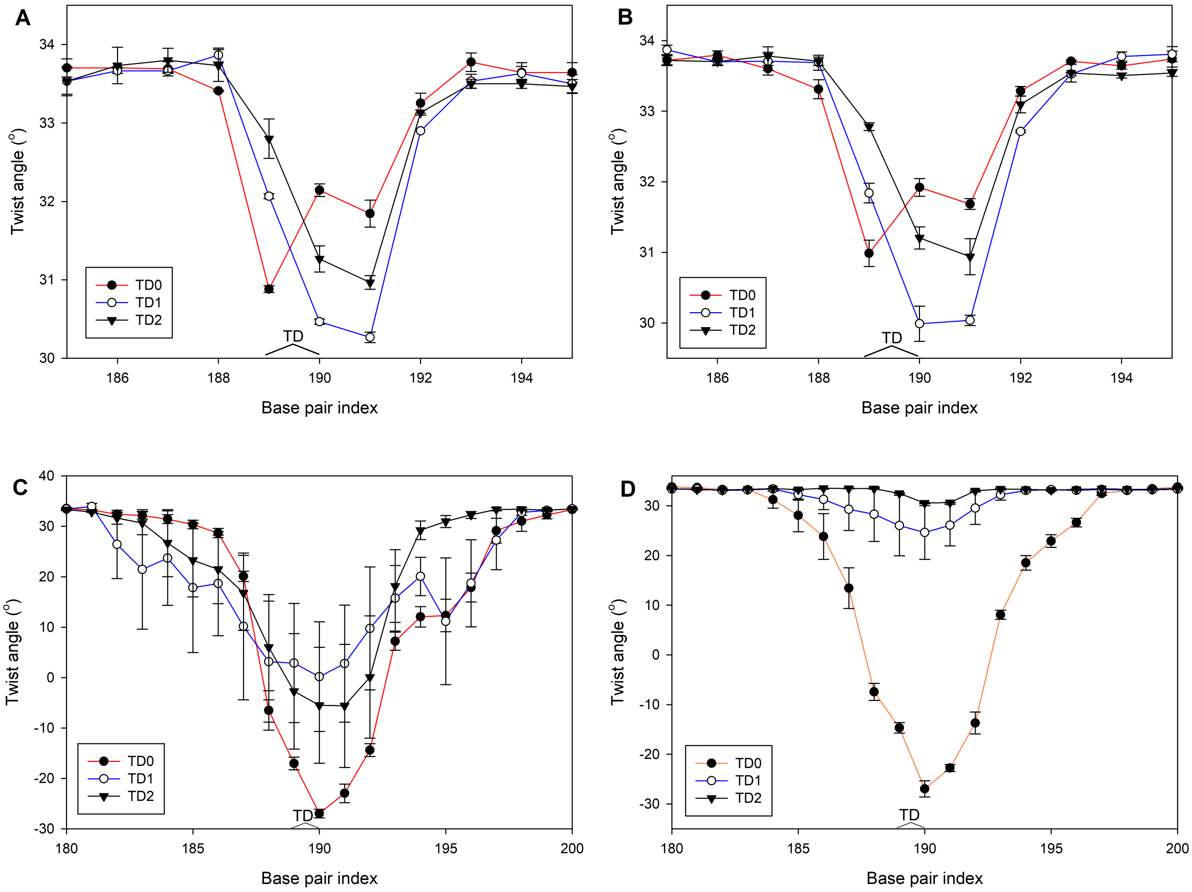


Figure S9: Twist angles measurements for the thymine dimer-containing dsDNA at state points (**A**) $F = 0.5$ pN and $\sigma \approx -0.021$ (B-DNA), (**B**) $F = 0.5$ pN and $\sigma \approx -0.056$ (plectoneme), (**C**) $F = 1.5$ pN and $\sigma \approx -0.073$ (tip-bubble plectoneme), and (**D**) $F = 2.0$ pN and $\sigma \approx -0.056$ (bubble). The value indicated at base pair index $i$ represents the twist angle between base pair $i$ and base pair $i-1$. Error bars represent standard errors of the mean. The thymine dimer is located at positions 189 and 190 and is marked by "TD" in the plots. In the B-DNA and plectonemic state, the largest decrease in twist angle (approximately 3.5°) is registered by TD1. This order changes when we consider the tip-bubble plectoneme and bubble states formed at higher forces, where TD0 shows the largest change in twist angle, because it shows the highest propensity for the tip-bubble or bubble to be localized at the thymine dimer at these state points. One of the reasons that TD0 differs so much from TD1 and TD2 for this bubble dominated state point is that for the latter, twist-induced bubbles are more likely to form at other locations along the DNA strands. How this changes as a function of force and twist is something that will be investigated in the future.

### S8 Force-torsion state diagrams for the TD models

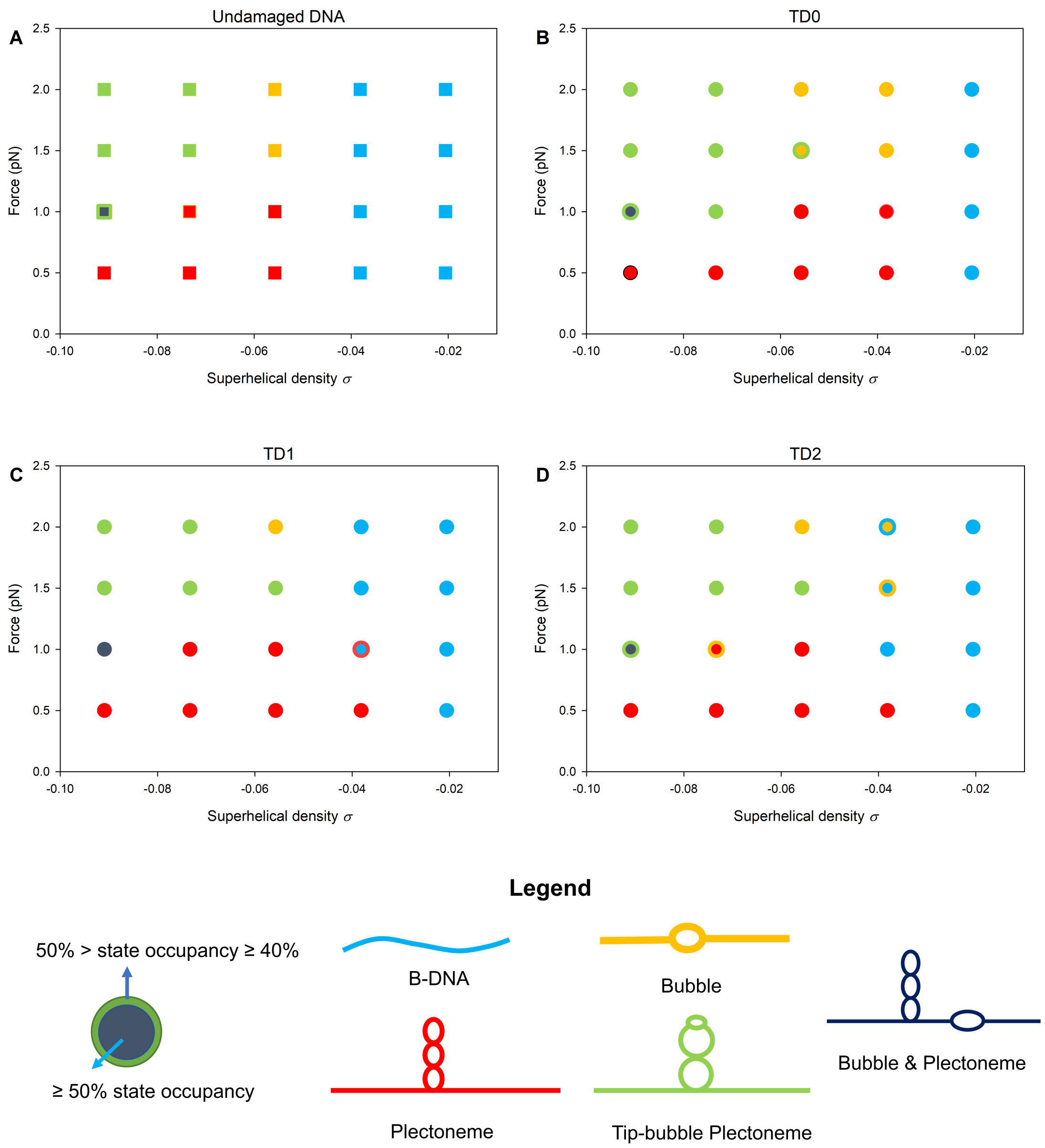


Figure S10: Force-torsion state diagrams for (**A**) undamaged and (**B** - **D**) thymine dimer-containing dsDNA. All three thymine dimer models produce similar enhancements in the state occupancy of plectoneme and plectonemic tip-bubble at lower superhelical densities (relative to the undamaged dsDNA).

#### S9 Plectonemic pinning for the TD models

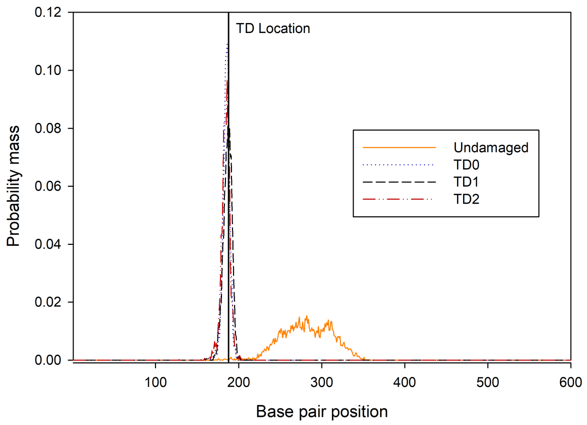


Figure S11: Plectoneme position distributions for a negatively supercoiled DNA strand at $F = 1.5$ pN and $\sigma = -0.073$. All three thymine dimer models yield very similar distributions.

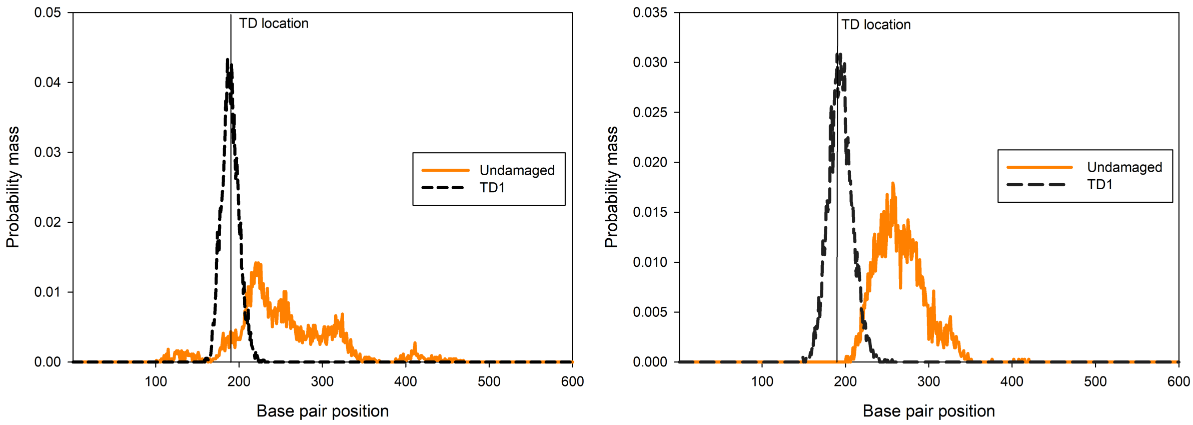


Figure S12: (Left) Plectoneme position distribution for a positively supercoiled DNA strand with TD1 at $F = 1.5$ pN and $\sigma = 0.067$. (Right) Plectoneme position distribution for a negatively supercoiled DNA strand with TD1 at $F = 1$ pN and $\sigma =$ -0.056. The combination of $\sigma$ and $F$ here ensures a relatively low tip-bubble state occupancy ($< 40\%$) for the trajectories. Under these state conditions, tip-bubbles are no longer the dominant state in the force-torsion state space, and therefore any observed plectonemic pinning can be directly attributed to the defect itself.

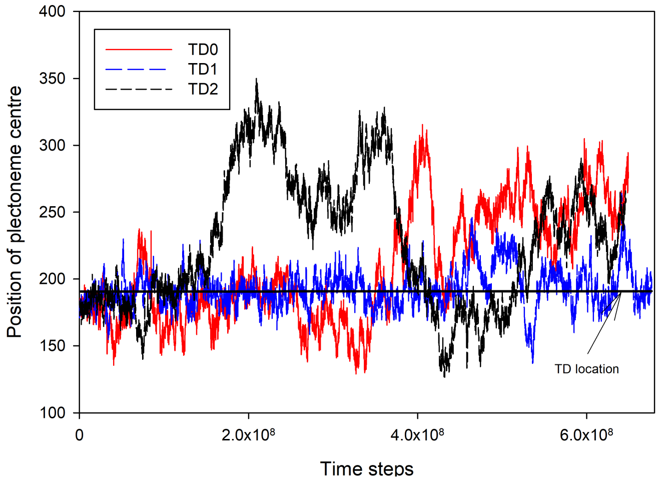


Figure S13: Pinning for positive supercoiling: Plectoneme centres are determined using the algorithm described in the supplementary material of Matek *et al.* [4]. With the exception of the choice of the thymine dimer model, other initial conditions (i.e. starting configuration, etc.) for the simulations are identical. TD1 is observed to pin the plectoneme more effectively at the chosen state point of $F = 1.5$ pN and $\sigma = 0.067$ than TD0 or TD2.

### S10 Denaturation probability of thymine dimer base pair

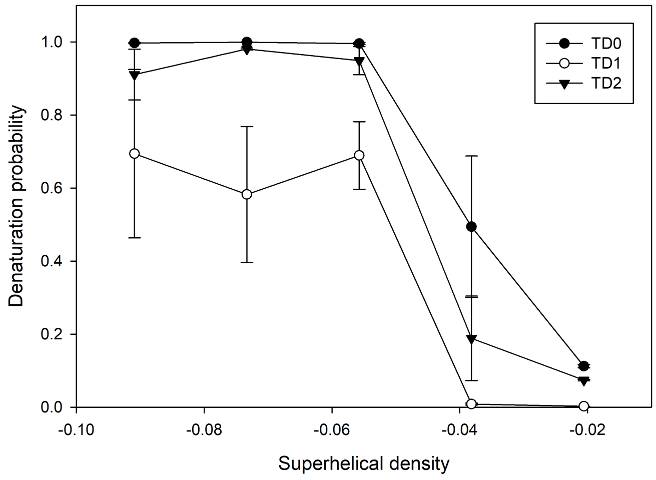


Figure S14: Denaturation probability of thymine-dimer base pairs at $F = 1.5$ pN, with error bars corresponding to standard errors of the mean. All three TD models show a marked increase in denaturation probability at the thymine-dimer base pairs when $|\sigma| > 0.04$. The localisation of the tip-bubble at the thymine dimer causes the thymine dimer to be denatured most of the time.

### S11 The structural effect of stacking interaction multiplier in thymine dimer models

The stacking of the thymine bases can be quantified indirectly by measuring the relative orientation angle of the thymine nucleotides as defined using their backbone-base vectors (Figure S15 for the definition).

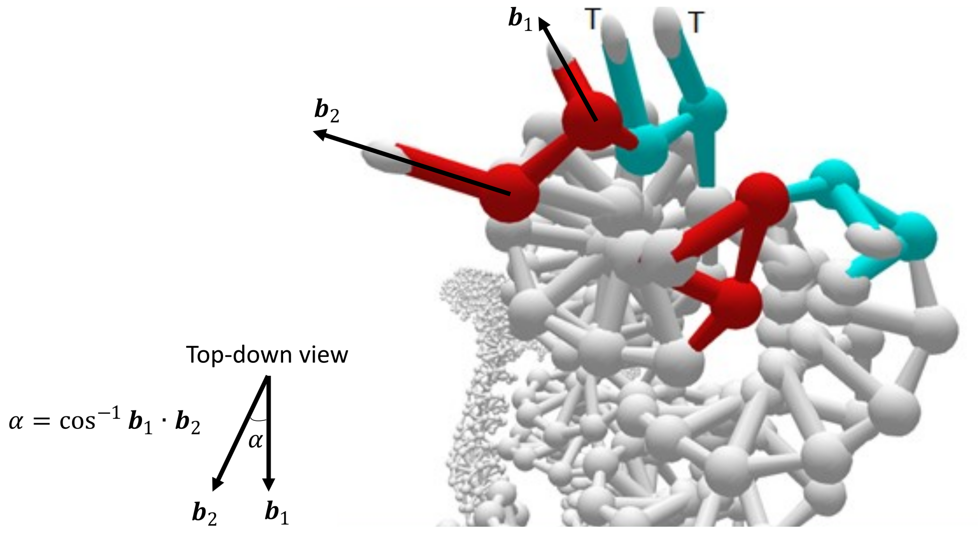


Figure S15: The relative orientation of two adjacent bases (in red) is defined by the orientation angle $\alpha$. $\hat{\mathbf{b}}_1$ and $\hat{\mathbf{b}}_2$ are the backbone-base vectors of the bases. The same definition applies for the thymine bases (labelled as T and coloured in cyan).

To illustrate that the thymine bases in TD0 remain stacked when in an extrahelical arrangement within a tip-bubble, orientation angles between the TD0 thymine bases were extracted from equilibration runs of supercoiled TD0 dsDNA at $F = 1.5\,\text{pN}$ and $\sigma = -0.073$. These conditions were chosen to ensure that the supercoiled dsDNA remained in the plectoneme tip-bubble state. For comparison, the relative orientation of a neighbouring set of bases in the tip-bubble was also measured, and the same approach was repeated for TD1 and TD2.

Figures S16 and S17 provide a clear contrast between the thymine bases in TD0 and those of TD1. With an enhanced stacking interaction, the thymine bases in TD0 remain stacked throughout the equilibration run, as would also be expected for a covalently bonded thymine dimer. Conformational changes associated with the formation of a plectoneme tip-bubble have no impact on the stacking between the thymine bases in TD0. On the other hand, the thymine bases in TD1 lose their stacking when the thymine dimer adopts an extrahelical arrangement within a tip-bubble. The same observation is true for TD2. This behaviour is a weakness of these two alternative models. Nevertheless, the global thermodynamic and pinning behaviour of these models is quite similar to TD0, suggesting that these particular properties are robust to variations in the stacking behaviour of the thymine dimer.

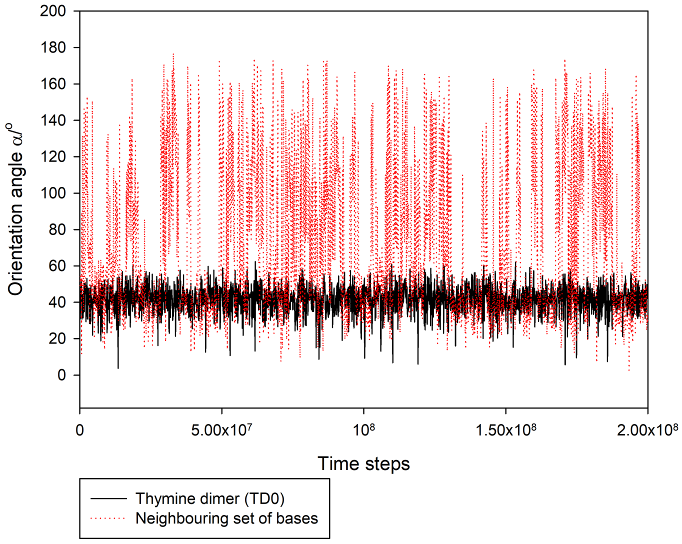


Figure S16: The orientation angles between the thymine bases in TD0 remain stable throughout a simulation run at $F = 1.5\,\mathrm{pN}$ and $\sigma = -0.073$ that is initiated with a tip-bubble plectoneme localized at the thymine dimer, whereas significant fluctuations in orientation are observed in the adjacent set of bases. This confirms that the enhanced stacking interaction multiplier in TD0 captures this expected consequence of the covalent bonding between the thymines.

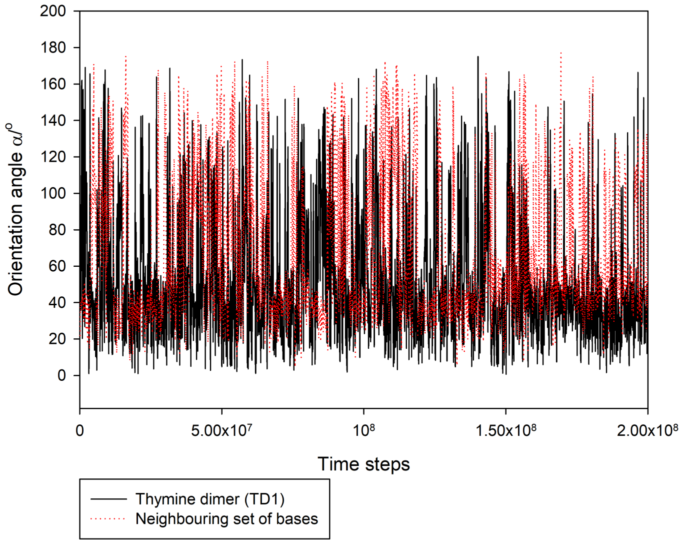


Figure S17: Without any enhancement of the stacking interaction in TD1, the thymine bases of TD1 frequently lose their stacking when the thymine dimer adopts an extrahelical arrangement within a tip-bubble at $F = 1.5\,\mathrm{pN}$ and $\sigma =$ -0.073. A similar behaviour is observed for TD2.
